# GōMartini 3: From large conformational changes in proteins to environmental bias corrections

**DOI:** 10.1101/2024.04.15.589479

**Authors:** Paulo C. T. Souza, Luís Borges-Araújo, Chris Brasnett, Rodrigo A. Moreira, Fabian Grünewald, Peter Park, Liguo Wang, Hafez Razmazma, Ana C. Borges-Araújo, Luis Fernando Cofas-Vargas, Luca Monticelli, Raúl Mera-Adasme, Manuel N. Melo, Sangwook Wu, Siewert J. Marrink, Adolfo B. Poma, Sebastian Thallmair

**Author notes:** Authors for correspondence: Paulo C. T. Souza. Adolfo B. Poma, Siewert J. Marrink, Sebastian Thallmair.

## Abstract

Coarse-grained modeling has become an important tool to supplement experimental measurements, allowing access to spatio-temporal scales beyond all-atom based approaches. The GōMartini model combines structure- and physics-based coarse-grained approaches, balancing computational efficiency and accurate representation of protein dynamics with the capabilities of studying proteins in different biological environments. This paper introduces an enhanced GōMartini model, which combines a virtual-site implementation of Gō models with Martini 3. The implementation has been extensively tested by the community since the release of the new version of Martini. This work demonstrates the capabilities of the model in diverse case studies, ranging from protein-membrane binding to protein-ligand interactions and AFM force profile calculations. The model is also versatile, as it can address recent inaccuracies reported in the Martini protein model. Lastly, the paper discusses the advantages, limitations, and future perspectives of the Martini 3 protein model and its combination with Gō models.

## 1. INTRODUCTION

The understanding of how proteins fold and perform their functions selectively, efficiently, and modulated by interactions with other biomolecules depends on the knowledge of their structure and dynamics. Despite tremendous progress in the experimental field^1,2^, molecular modeling techniques have conquered their own space as an important and complementary set of approaches to study proteins^2,3^. In particular, given the limitations in obtaining experimental high-resolution atomistic details from short to long time-scales, all-atom (AA) molecular dynamics (MD) simulations have become widely used to study protein dynamics and even folding of simple systems^4,5^. Because of the high computational costs associated with AA MD, these are usually limited to studying phenomena occurring on time scales of 1-100 µs (depending on the protein and/or system size)^6,7^. Thus, a broad range of important biological phenomena remains out of reach. For instance, these include the long-range motion of protein domains as well as fit-induced mechanisms involving protein-ligand or protein-protein interactions. Systems involving transmembrane or peripheral membrane proteins can be even more challenging, as protein dynamics in a lipid bilayer environment can be slowed down, coupled to membrane fluctuations, and possibly dependent on lipid composition^8,9^. AA approaches are also heavily challenged by the interpretation of experimental data such as single-molecule force spectroscopy by atomic force microscopy (AFM-SMFS) profiles generated in nanomechanical studies, where it is critical to accumulate enough sampling of non-equilibrium pulling processes^10–12^. Similarly, in the case of disordered proteins or domains, integrating small-angle x-ray scattering (SAXS) with MD data requires the determination of whole ensembles of conformations which can be difficult to obtain with AA methods^13,14^. Enhanced sampling methods and GPU parallel computing help to reach longer timescales and better sampling with AA approaches^15–17^. However, when considering biological length- and timescales, they are still limited to rather local processes.

One attractive alternative to AA protein models is the use of coarse-grained (CG) approaches. CG models are simplified representations of AA models which, due to a reduction of explicit degrees of freedom and a smoother interaction landscape, offer a substantial simulation speed up. As a result, CG models can reach length- and timescales which are orders of magnitude larger than AA models. CG approaches offer a wide range of resolutions and strategies to define their interactions^18–20^. For instance, structure-based protein models, like Gō-type models^21–23^, define their interactions based on a known and usually folded structure. The potential energy (*U*_*G*_*_ō_*) in the Gō model for proteins is constructed based on a well-defined native structure of the protein with a functional form as follows, *U*_*G*_*_ō_* = ∑^*NC*^_i<j_ *V*(σ_*ij*_, ε), where NC denotes the set of native contacts in the protein structure and *V*(σ_*ij*_, ε) is a Lennard-Jones (LJ) 12-6 potential. σ_*ij*_ is given for each native contact and depends on the distance between specific pairs (i.e. σ_*ij*_ = *r*_*ij*_/*2*^1/*6*^). *V*(σ_*ij*_, ε) is parameterized based on the folded structure. The ε represent the energy scale of the native contacts and is usually uniform for all contacts. Typically representing each residue as a unique particle, Gō-like approaches are a useful tool for modeling near-native protein dynamics. However, environmental effects are usually neglected^18^. On the contrary, physics-based models, such as the well-known Martini force field^24–26^, can be used to model protein dimerization and aggregation processes as well as interactions with lipid bilayers and other biomolecules^8,9,18,27^. With each protein residue being represented by 1-5 beads, Martini still retains chemical specificity, because the beads are parametrized using experimental thermodynamic data such as partitioning free energies of small compounds between polar and apolar environments. While bonded potentials are parametrized and validated using atomistic and experimental data^24–27^, traditional Martini protein models exhibit limitations in accurately representing stably folded proteins, often relying on a harmonic elastic network to maintain structural stability^28^. Although the dynamic accuracy of elastic networks can be improved via neural network-based structure predictions^29^, extensive tests have also shown the combination of Martini with elastic networks may also contribute to inaccurate protein−protein interactions^28,30^. The observed stickiness of proteins^31–33^ in Martini 2 may also affect their accessible conformational ensemble.

A possible way to keep a good compromise between high computational performance, accurate protein dynamics, and reliable interactions with the environment is the combination of structure-based and physics-based coarse-grained approaches. A recent example for this is the combination of Gō and Martini 2 models, called GōMartini^34^. Several studies have shown that GōMartini models can be parametrized to reproduce protein flexibility from atomistic benchmark simulations^34–36^. In addition, GōMartini has also shown great potential to study the nanomechanical stability of proteins^30,37^. However, it has limitations in reproducing longer-range conformational changes^34^. In addition, the model inherited parts of the stickiness limitations of Martini 2^30^.

Here, we present the virtual-site implementation of an enhanced GoMartini model, which can be combined with the latest iteration of Martini, together with a diverse set of applications and comparisons to elastic network models. Moreover, the fully reparameterized Martini 3 model for proteins is presented, pointing to which improvements in the model may enable more accurate predictions of protein packing and protein interactions^38,39^. The GōMartini implementation together with Martini 3 has already shown that it can capture subtle changes in protein dynamics caused by interactions with membranes^40^, single point mutations^41^ and mechanostability.^42^ We also show how the virtual-site implementation can be used to implement an environmental bias to correct recently described inaccuracies of the model, such as underestimated dimensions of intrinsically disordered proteins (IDPs)^43,44^ and low hydrophobicity of certain amphiphilic small peptides.^45^ The paper is structured as follows: first, we discuss the Martini 3 protein model followed by the changes in the enhanced GōMartini model as well as the improved virtual-site implementation facilitating high parallelization. We further demonstrate the power of the GōMartini model using four case studies: (i) binding of a Pleckstrin homology (PH) domain to PI(4,5)P2-enriched membranes, (ii) binding of benzene to T4 lysozyme, (iii) an allosteric pathway in Cu,Zn superoxide dismutase, and (iv) AFM-SMFS force profile calculations for the case of protein complexes such as antigen:antibody and dockerin:cohesin systems. Next, we give a perspective on how GōMartini models can be further optimized and moreover, how virtual Gō particles can be used to introduce environmental bias corrections through changes in the interaction with water beads. In the final section, we discuss the advantages, limitations, and future prospects of the approach and the overall development of the Martini 3 protein model.

## 2. THE MARTINI 3 PROTEIN MODEL

The Martini 3 protein model is the natural evolution of the previous Martini 2 iteration^26^, which now leverages the improvements introduced with the Martini 3 force field. However, it can still be considered as a prototype model, just like the current Martini 3 lipids, since the model has not been fully updated with the current parametrization rules. In particular, the core of the protein model is still based on a single particle backbone (BB), which is placed at the center of mass (and not the center of geometry) of the N, Cα, Cβ and O atoms of the underlying atomistic backbone, and to which 1-5 side chain (SC) beads may be attached. This is in line with the original implementation of the Martini protein force field^25^, and differs from the ELNEDIN model^28^, where the BB bead was placed at the position of the C*α* atoms. To connect consecutive amino acid residues, a harmonic bond or constraint, depending on the secondary structure, is placed between their BB beads. Angle and dihedral potentials are then placed over 3 or 4 consecutive BB beads, respectively, to define the secondary structure-dependent backbone torsion behavior. This set of BB bonded parameters – composed of the bond lengths, angles, dihedral angles, and their respective force constants for each of the secondary structure motifs – was inherited from the original implementation of the Martini protein force field^25^. These were parameterized from a representative set of ∼2000 proteins from the protein data bank (PDB), on which the Define Secondary Structure of Proteins (DSSP) algorithm^46^ was used to determine the secondary structure motif associated with each residue.

In the original implementation of the Martini protein force field, the BB particle type depended on the secondary structure motif associated with a respective residue^25^. When free in solution or in a coil or bend it was represented by a P5 bead; BB particles in beta strands or turns were represented by Nda beads; and in helices by N0 beads, with the C- and N-termini of a helix represented by Na and Nd beads, respectively. This choice was made to better represent the inter-backbone hydrogen bond character of each residue when present in a specific motif — i.e. the hydrogen bonds established within a helix would reduce the polar character of the amide backbone group^25^. In contrast thereto, all backbone beads are represented by P2 beads in Martini 3, regardless of the underlying secondary structure motif. The exceptions to this rule are charged terminal backbone beads — which are represented by Q5 beads — and GLY, ALA, VAL, and PRO residues. These four amino acids use different bead types to better represent slight differences in chemical group polarity and size. The GLY backbone is mapped as an SP1 bead to represent the loss of the side chain (but keeping similar polarity compared to the default P2 backbone), while the PRO backbone is mapped as an SP2a, due to the lack of hydrogen-donor capabilities. ALA and VAL are mapped as SP2 beads, to avoid overmapping issues which could be caused by their side chain particles being mapped quite close to the backbone^30^.

The side chains have been completely revisited for the Martini 3 protein model, following the new parameterization guidelines established with the Martini 3 release and making use of the larger number of bead types and new bead sizes specific for mappings finer than 4-to-1^39^. The side chain models were parameterized from their backbone-less analogues, and calibrated considering their molecular volume, partitioning behavior, solvent properties, and miscibility trends. The mapping and bead assignments of the Martini 3 protein model are shown in Figure 1. The parameterization of the side chain analogues is described in detail elsewhere^39,47^. To illustrate the quality of the current side-chain models, partitioning free energies of their analogues in three different water/oil systems are compared with experimental data in Table S1 and Figure S1.

**Figure 1.**
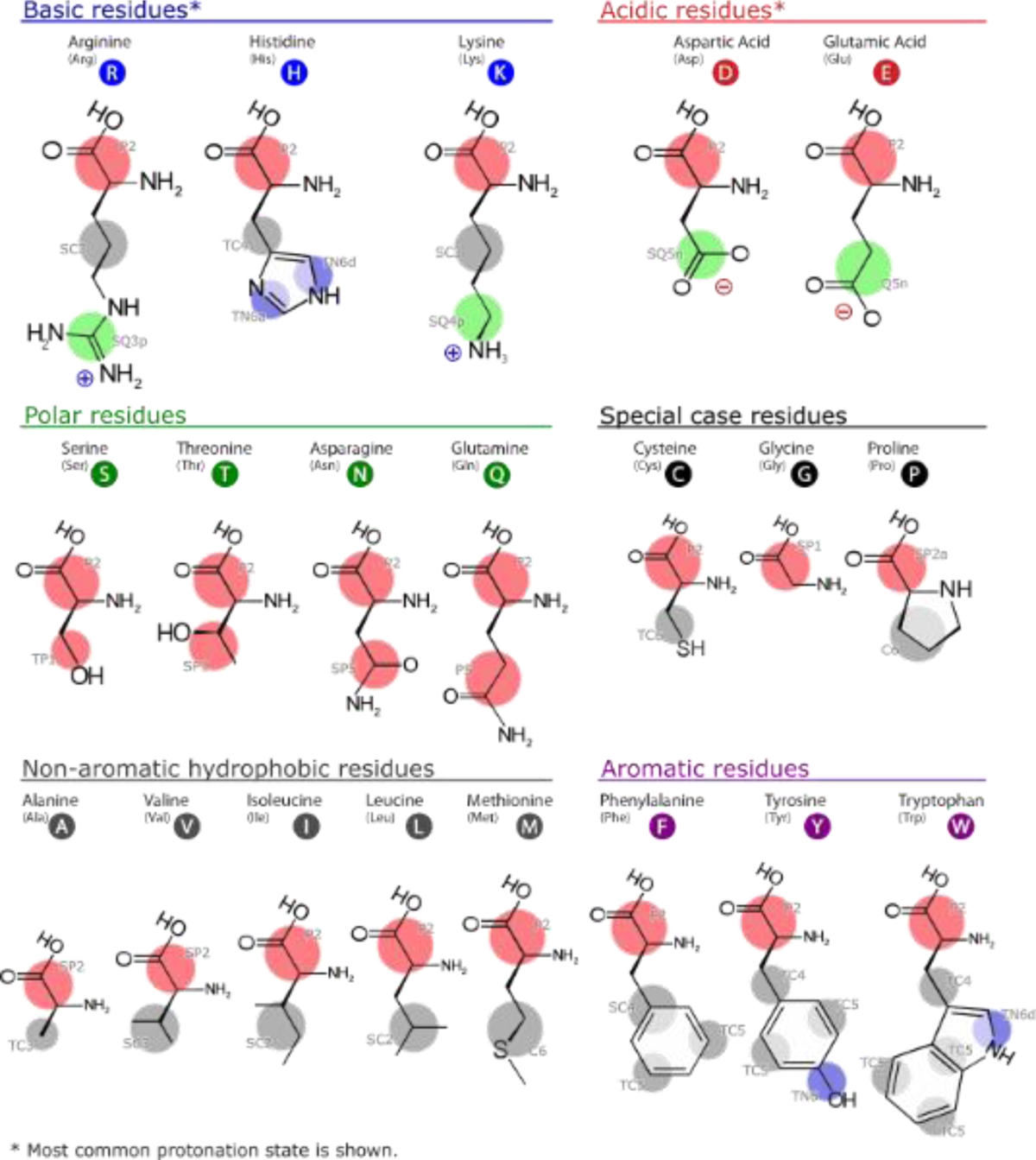
Mapping and bead chemical types of the Martini 3 protein model. The colors indicate the main classes of bead chemical types: P (polar, in red), N (intermediately polar, in blue), C (nonpolar, in gray) and Q (charged, ingreen). Different bead sizes are also indicated, ranging from the bead with the largest radii (regular, no symbol) to small (S) and tiny (T) beads.

Like in previous Martini protein models, the side chains are connected to the BB bead via harmonic bonds or linear constraints and their dynamics are controlled by two angles spanning the first SC bead, the BB bead, and the BB bead of the two neighboring residues (*-BB BB SC1* and +*BB BB SC1*). In the Martini 3 protein model, the use of the side chain dihedral corrections (side chain fix – scFix)^48^ becomes standard for protein models with defined secondary and tertiary structure. The scFix restrains the torsional flexibility of the side chains by adding dihedral potentials spanning the first SC and BB beads of consecutive residues (*SC1 BB +BB +SC1*, *SC1 BB +BB ++BB*, and *SC1 BB -BB --BB*) and thus preventing unrealistically high side chain flexibility^48^. The dihedral equilibrium angles are set at the value obtained from the atomistic reference structure used to build the Martini model.

The Martini 3 protein model still requires the use of a tertiary structure bias, such as an elastic network or Gō-like model, to maintain the native folded structure of proteins. The current elastic network implementation for Martini 3 applies harmonic bonds between BB beads based on a cutoff criterion, and with a single force constant for all bonds. Standard values of 0.85 nm and 700 kJ/(mol nm^2^) are recommended for the upper distance cutoff and force constant, respectively. The value of the recommended force constant was slightly increased in relation to 500 kJ/(mol nm^2^) commonly used in Martini 2, as in certain scenarios this value could be too low, inducing an increased level of stickiness^30^. Apart from the distance cutoff, a residue pair must be separated by at least two residues for an elastic network bond to be applied between them. For instance, residue *i* can be bound to residue *i*+3, which corresponds to a sequence distance of *k* = 3. While the current elastic network successfully maintains folded structures, it also prohibits studies which may involve conformational changes or unfolding, due to the unbreakable harmonic bonds which are used to build the network.

Martini 3 protein models have been validated against a large variety of systems, including test cases reported in the main publication^39^ and a series of spin-off studies published in separate works^38,49–52^. Examples of performed validations are: aggregation levels of soluble proteins in water and polyleucine helices in lipid bilayers, di^53^merization free energies of transmembrane (TM) peptides, binding of ions^39^ and small molecules to proteins^38^, biomolecular condensates in different ion concentrations^53^, and lipid interactions with transmembrane and peripheral proteins^49-52^.

## 3. NEW IMPLEMENTATION OF THE GŌMARTINI MODEL

### Virtual-site implementation

The new implementation of the Martini Gō-like model relies on the use of virtual interaction sites, which are constructed using the position of the BB bead as reference. The virtual interaction sites are solely used to define the interactions within the Gō-like model, which are encoded as Lennard-Jones (LJ) potentials between virtual site pairs. By default, these particles do not interact with any other beads in the system. As an extended feature, the use of virtual sites allows changing specific interactions between proteins (BB particles) and other beads in the Martini interaction table without compromising the integrity of the original force field. For example, we show that a secondary structure-specific water bias can be applied to improve properties of IDPs.

The major advantage of using virtual interaction sites is that it enables the use of non-bonded cutoffs as implemented in GROMACS. In the original 2017 implementation of the GōMartini model^34^, the LJ potentials were defined in GROMACS as pair potentials within the protein topology, which are treated internally as bonded potentials. Consequently, no cutoff was applied to the potentials. This is not problematic as long as the minimum of the potential is close to or below 1 nm, and the distance of the connected beads stays in the region of this minimum position. However, because the GōMartini model aims to allow for more conformational flexibility — including the dissociation of some of the native contacts — the lack of a cutoff can severely restrict the applicability of the model. One of these restrictions was the incompatibility of the original implementation with increasing parallelization due to specificities of the domain decomposition implementation in GROMACS. In practice, the parallelization of a simulation with a moderately-sized transmembrane protein, such as the light-harvesting complex II, embedded in a small membrane patch with a system size of ∼19.700 CG beads^35^ was restricted to about 10 processors. Our implementation based on virtual interaction sites circumvents this limitation at the minor cost of describing the BB of each amino acid by two CG beads instead of one. Considering the overall number of CG beads present in a typical system, the number of BB beads is usually only a minor fraction of a few percent. Thus, doubling the number of BB beads only slightly increases the total number of CG beads in the system.

### Enhanced GōMartini model

Besides the improved implementation, a few additional features were also adapted. While the contact map calculation remains unaltered from the original implementation – defined by residue overlap (OV) and restricted chemical structural units (rCSU) criteria^34^ – contacts in the contact map are now only included in the GōMartini model, if they are within a certain distance range in the reference structure. We used a range between 0.3 – 1.1 nm. The lower boundary was chosen to avoid regions with excessively high bead density. These can create artifacts due to increased interactions with the surrounding, especially with other high bead density regions^30^. The upper boundary was set to the non-bonded cutoff used in Martini simulations. Thus, only contacts, which have their minimum position within the non-bonded cutoff, are included in the model. Note that the underlying distance is measured between the BB beads of the Martini protein model. Thus, contacts are rarely excluded based on the lower boundary, while a few contacts are usually excluded due to the upper boundary.

The minimum sequence distance of the original model is *k =* 3.^34^ In our implementation, we used a minimum graph distance of *k =* 4, since at *k =* 3 the relative positions of BB particles can still be largely defined by bonded terms. Note that in the graph distance space of *Martinize2* – the tool for automatic Martini protein topology generation^54^ where we implemented the enhanced GōMartini model –, are not only sequential BB–BB bonds considered, but also disulfide bridges. In any case, we recommend *k =* 3 in cases where the protein flexibility in loops is too high because no dihedrals are defined there.

Furthermore, regular non-bonded interactions – i.e. the Martini bead-bead interactions – between pairs of BB beads are excluded in the enhanced GōMartini model if the amino acids have a contact according to the contact map. The reason for this choice is that the minimum position *rmin* of the sum of two LJ potentials, one from GōMartini and another one from the regular non-bonded Martini 3 interactions, is effectively at the larger *rmin* of the individual LJ potentials if the depths (*ε*) of the potentials are comparable. As long as the distance between the BB beads in the reference structure is larger than the *rmin* of the regular non-bonded LJ potential, this does not impact the protein structure. However, if the reference structure has a shorter *rmin* the protein structure gets distorted. Excluding regular non-bonded interactions between BB beads connected in the GōMartini model avoids this distortion. Overall, adding these exclusions is an upgrade relative to the previous GōMartini implementation, as it improves protein packing in regions involving backbone-backbone interactions, such as beta-sheets.

### General workflow to set up the GōMartini model

In order to build a GōMartini model for a protein, a two-step procedure has to be followed. First, a contact map specifying the OV and rCSU contacts can be obtained from the web-server http://pomalab.ippt.pan.pl/GoContactMap/^36,37^ (which is replacing the previous one^55^: http://info.ifpan.edu.pl/~rcsu/rcsu/) or via the ContactMapGenerator program available at https://github.com/Martini-Force-Field-Initiative/GoMartini/tree/main/ContactMapGenerator, using the default settings^55,56^. Subsequently, *Martinize2* can be used to obtain the CG coordinates and topology files from the atomistic reference structure and the contact map.^57^ Figure 2 summarizes this workflow to set up the GōMartini models. In order to activate the GōMartini model for the structure bias in *Martinize2*, the ‘-go’ flag has to be provided in addition to providing the contact map file using the ‘-go-map’ flag. Note that the format of the contact map has to adhere to the specifications outlined in the Supporting Notes B2. The Gō model can further be fine-tuned by adjusting the biasing strength (‘-go-eps’), the upper and lower cut-off distance (‘-go-up’ and ‘-go-low’), as well as the residue distance (‘-go-res-dist’). If these flags are omitted, *Martinize2* uses the default values described in the previous section. The default value for the potential depth is still the one recommended in the original GōMartini implementation: *ε*LJ = 9.414 kJ/mol. In contrast to the previous implementation, *Martinize2* utilizes the graph residue distance instead of the sequence distance, which are the same for almost all cases except those where amino-acids are connected through the side-chains (e.g. in the case of disulfide bridges). Aside from the definition of the intra molecule parameters, *Martinize2* writes the atom types and non-bonded interactions required to run a simulation with the GōMartini model. The file name of these files is preceded with the molecule name (by default ‘molecule_0’), which is also used in the naming of the virtual sites. Utilizing the ‘-go-moltype’ flag the name can be adjusted, such that when martinizing multiple different proteins the Gō definitions are compatible. More details including an example and tips and tricks are provided on the *Martinize2* github page (https://github.com/marrink-lab/vermouth-martinize). The enhanced GōMartini using virtual sites is also implemented in MAD, the Martini Database^58^, which also includes the option to manually remove or add Gō interactions, allowing the user to include for instance additional experimental information which may correct issues originating from the reference structure.

**Figure 2:**
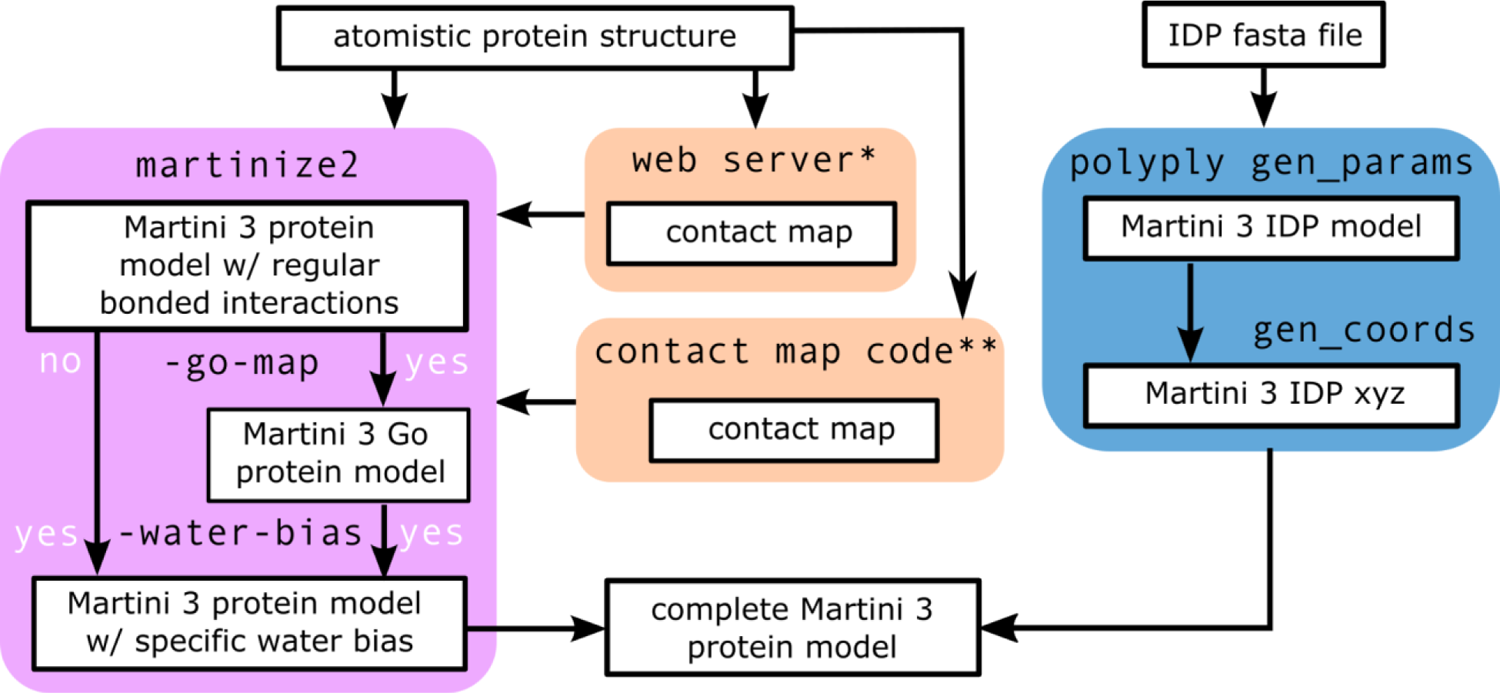
Workflow to set up the GōMartini model with contact maps for structured proteins and corrections of the BB-water interactions, respectively. Contact maps can be generated by a web-server http://pomalab.ippt.pan.pl/GoContactMap/^36,37^ or via the contact map program available at https://github.com/Martini-Force-Field-Initiative/GoMartini.

### Adding virtual site water bias

As previously mentioned, the virtual sites approach underlying the enhanced GōMartini model can also be used to specifically fine-tune interactions between Martini beads. For instance, it can be used to modify the strength of the protein-water interaction. Using the ‘-water-bias’ flag in *Martinize2* allows to automatically generate the BB virtual sites and non-bonded interaction parameters required for the water bias. Whereas the water bias can be combined with the GōMartini model, it does not require it. For example, adding water bias with an elastic network is also possible. The water bias can be added depending on the secondary structure element, with its strength defined using the ‘-water-bias-eps’ flag. The water bias defines the depth of the LJ potential between the virtual site and the water bead. The values can be positive (to effectively increase the water-BB virtual site interactions) or negative (decreasing water-BB virtual site interaction). The default value for strength of the water bias in *Martinize2* is zero, so the user needs to define it. Some suggestions are presented in Section 6. Finally, *Martinize2* also supports the definition of intrinsically disordered regions (IDRs) using the ‘-id-regions’ flag. A special water bias can be defined for these regions with the previous flag using ‘idr’ instead of a secondary structure assignment. Defining IDRs is useful when a protein contains both a folded and disordered domain, because *Martinize2* might still detect some (transient) secondary structure in the disordered domain.

### Martini 3 IDPs from sequence

Lastly, Martini 3 CG protein models for IDPs can also be generated directly from a sequence fasta file using *Polyply*^59^. As a fully disordered protein has no reference structure, a contact map is not necessary for this method of generation. Instead *Polyply* gen_params automatically generates the molecule itp file from the sequence already including the virtual sites for the addition of an appropriate water bias, and the automatic addition of additional backbone dihedral potentials. As *Polyply* should not be generally used to generate topologies of folded proteins, these parameters are added automatically (ie. without any additional flag specifications) when *Polyply* is used with the Martini 3.0.0 force field library. System coordinates (e.g. of a solvated IDP) can subsequently be generated using *Polyply* gen_coords (Figure 2). The *Polyply*-generated files can be used directly, only externally requiring a further definition of the extra interactions between the BB virtual sites and water as in the case of *Martinize2* and described in more detail in the previous section. An example of how to use *Polyply* to generate and use these parameters is available on the *<u>Polyply</u>* <u>github wiki</u>. The *Polyply* route is especially convenient for high-throughput simulations of many different disordered proteins.

## 4. SIMULATION DETAILS

### General simulation settings

All simulations were performed with the program package GROMACS^60^ (versions 2018.x to version 2023.x). Settings for the CG simulations follow the “new” set of Martini run parameters^61^ using a time step of 20 fs. Specifically, the Verlet neighbor search algorithm was used to update the neighbor list, with a cutoff of 1.1 nm for the non-bonded interactions. Coulombic interactions were treated using reaction-field electrostatics with a dielectric constant of 15. The Parrinello–Rahman barostat^62^ (coupling parameter of 12.0 ps) and the velocity-rescaling thermostat^63^ (coupling parameter of 1.0 ps) were used to maintain pressure and temperature, respectively. More technical details about the system setups, simulation settings, and analysis for each specific test case are given in the Supplementary Methods.

### Martini models and system setup

All simulations were performed using the open-beta^64^ or more recent development versions of Martini 3 force field^39^, with the protein models generated by *Martinize*^26^ (for open-beta test cases), *Martinize2*^54^ (for folded proteins simulated with the final Martini 3 release) or *Polyply*^59^ (for IDPs simulated with the final Martini 3 release). Except for IDP and biomolecular condensate systems, bonded parameters are still dependent on the secondary structure, which is calculated by the DSSP approach^46^ using an atomistic reference structure. In addition, the side-chain dihedral corrections scFix^48^ are included for all secondary structure elements. The contact maps for the GōMartini models were generated using the contact map approach proposed by Cieplak and co-workers^55^ (http://info.ifpan.edu.pl/~rcsu/rcsu/) or the new implementation from the Poma group^36,37^ (http://pomalab.ippt.pan.pl/GoContactMap/). Besides the new implementation of the GōMartini model, we also employed two different elastic network (EN) models to maintain the structural protein scaffold in the case of the PH domain, T4 lysozyme, and SOD1. Slightly modified versions of these two EN settings are typically used in combination with the CG force field Martini^25,26,28,30^. Here, we want to focus solely on the impact of the structural bias models. Thus, all other bonded parameters of the protein models are unchanged between the Gō-like and the EN models. The EN models are set up based on a distance cutoff criterion between the BB beads in the CG reference structure of the protein. Harmonic potentials are used to constrain the protein flexibility and to maintain the protein structure. These can be applied in two different ways: (i) either the non-bonded interactions between the BB beads connected by a harmonic potential are excluded or (ii) the harmonic potentials act on top of the non-bonded interactions. In the first case, the bond type corresponds to a regular chemical bond, i.e. bond type 1 in GROMACS. In the second case, bond type 6 is used in GROMACS. In the following, we use the GROMACS bond types to distinguish between the different settings, namely EN type 1 and EN type 6, respectively. See more details about the protein models used in each specific test case in the Supplementary Methods.

## 5. CASE STUDIES: SHOWING THE ADVANTAGES OF GŌMARTINI 3

### PLCδ1 PH domain: PMFs for strong lipid binding

In the first case study, we investigated the membrane binding affinity of the PH domain of the phospholipase C PLCδ1. It is a peripheral membrane protein and a representative of the phosphoinositol phosphate (PIP) binding family of PH domains. The PLCδ1 PH domain discussed here favorably binds PI(4,5)P2 ^65–67^. Figure 3A shows the PCLδ1 PH domain with a PI(4,5)P2 lipid in its crystal structure binding pocket embedded in a POPC bilayer. Unbiased MD simulations starting from the membrane-bound structure confirm the high affinity of the PLCδ1 PH domain to a single PI(4,5)P2 lipid (Figure S2) which has also been shown previously based on atomistic as well as CG simulations^40,49,65,67,68^. We used three different structural bias models. Besides the GōMartini model, two different EN models with a cutoff distance of 0.8 nm were used. One was described by bonds of type 6 and a force constant of 500 kJ/(mol nm^2^), hereafter EN6, while the other one had bonds of type 1 and a force constant 700 kJ/(mol nm^2^) (hereafter EN1; for details of the models see Section B1 of the Supporting Methods). All three models confirm the strong binding of the PLCδ1 PH domain to PI(4,5)P2. In the case of GōMartini and the EN1, the protein unbinds in one replica each but it is able to find the PI(4,5)P2 lipid again and re-binds to it (Figure S2).

**Figure 3.**
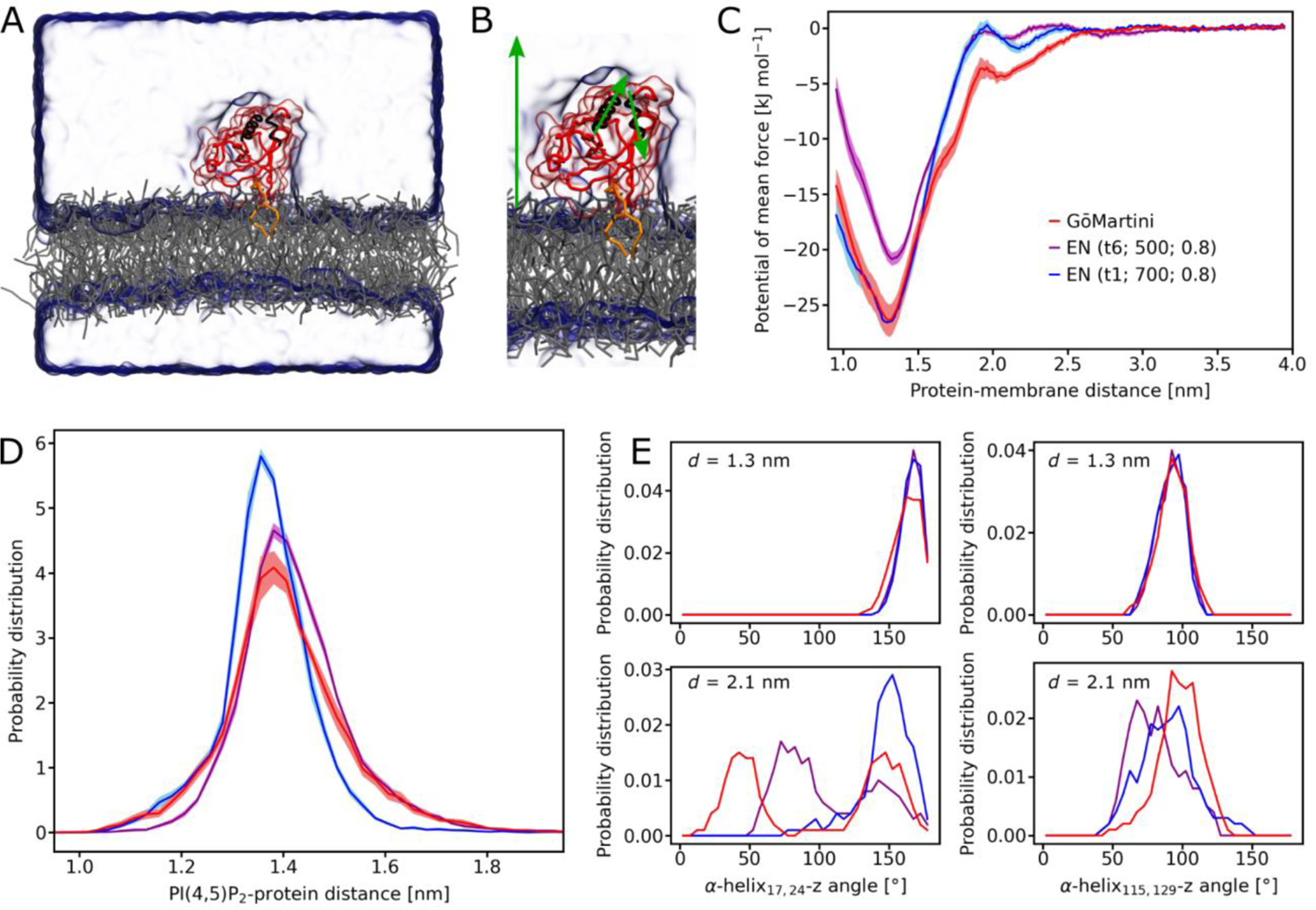
PI(4,5)P2 binding of the PLCδ1 PH domain studied at the CG Martini level. A) Setup of the simulation box containing a POPC bilayer (grey) with one PI(4,5)P2 (orange) in the binding pocket of the PLCδ1 PH domain (red) solvated in water (light blue transparent surface). B) Magnified view of PLCδ1 PH domain with the green arrows indicating the vectors used to determine the orientation: membrane normal z (left), α-helix115,129 (middle), and α-helix15,24 (right). C) Potential of mean force for the PI(4,5)P2 binding of the PLCδ1 PH domain. The protein is modeled with the GōMartini model (red) as well as two different elastic network models of type 6, force constant of 500 kJ/(mol nm^2^), cutoff 0.8 nm (violet) and of type 1, force constant 700 kJ/(mol nm^2^), cutoff 0.8 nm (blue). D) Probability distribution of the protein-membrane distance evaluated for ten replicas of 2 µs for each protein model. The distance is measured between the PI(4,5)P2 head group and the center of mass of the protein. E) Orientation of the PLCδ1 PH domain measured by evaluating the probability distributions of the angles between α-helix15,24/α-helix115,129 and the membrane normal z. The colors in D) and E) are the same as in C).

In order to quantify the binding affinity of the PLCδ1 PH domain, we calculated the potential of mean force (PMF) for the protein-lipid unbinding following the protocol of Naughton *et al.*^69^. Figure 3C depicts the corresponding PMFs along the distance between the lipid head group and the protein center of mass obtained with the three structural bias models. All three models confirm the strong affinity of the PLCδ1 PH domain to the PI(4,5)P2 head group with a minimum at a protein-membrane distance of ≈1.30–1.33 nm. The GōMartini model exhibits a minimum distance of 1.31 nm while it is slightly shifted for the EN models (EN1: 1.30 nm, EN6: 1.33 nm). Overall, the three models agree on the location of the PMF minimum. The PMFs exhibit further differences between the three models. First, the error bars are larger in the case of GōMartini. This is expected because it contains fewer bonds than the EN models and because the LJ potential allows for more flexibility so that in extreme cases, contacts can dissociate completely. As this increases the accessible conformational space, more sampling time is required to achieve the same level of error. Second, the depth of the minimum differs between the models. While the PMFs of the GōMartini model and the EN1 model are the same within the error bars around the global minimum (minima at −26.3±1.4 kJ/mol and −26.7±0.9 kJ/mol, respectively), the PMF of the EN6 has the highest minimum at −21.1±0.6 kJ/mol. In order to better understand the effect of bond type (and exclusions) on binding affinity, an additional PMF calculation was performed using bond type 1 with an EN force constant of 500 kJ/(mol nm^2^) (instead of 700 kJ/(mol nm^2^)). Excluding LJ interactions (bond type 1) lowers the PMF minimum by about 15 kJ/(mol nm^2^) (Figure S3). This result confirms an observation reported recently that EN models can overestimate the aggregation between proteins if the force constant is too low (here 500 kJ/(mol nm^2^)) and the non-bonded interactions in the network are excluded^30^, due to a high bead density which can result in an overestimation of the interaction energy.

A characteristic of several PH domains is the existence of two binding modes to PIP lipids: a tightly bound structure corresponding to the crystal structure binding pocket and a loosely bound structure^65^. Two different PIP interaction sites are known for PH domains: the canonical C-site and an alternative A-site which is the less common binding site. For the tightly as well as the loosely bound structure both orientations have been detected^65^. The PLCδ1 PH domain studied here preferentially orients its C-site towards the membrane at shorter and longer PI(4,5)P2 protein distance^65^. Figure 3C shows that the three models differ also in the binding strength of the loosely bound structure. While the GōMartini model exhibits the highest stabilization, the EN models show a reduced stabilization by more than 50%. To better understand the changes in orientation between the tightly and loosely bound structures, we analyzed the angles between two α-helices - α-helix15,24 and α-helix115,129 - and the membrane normal z for two windows of the umbrella sampling depicted in Figure 3E. For the tightly bound structure, the probability distributions of the angles show a good agreement (upper panels, *d* = 1.3 nm). This changes at the loosely bound structure (lower panels, *d* = 2.1 nm). Here, the probability distributions of the α-helix15,24-z angle differs between the three models. The GōMartini model stabilizes two orientations at 45° and 145°. Also EN6 stabilizes two orientations (75° and 145°), while for EN only an orientation similar to the tightly bound structure is observed. This suggests that the GōMartini model allows the protein to better adjust to the loosely bound structure which stabilizes the interaction with the membrane.

### T4 lysozyme: small-molecule binding

Engineered mutants of T4 lysozyme are known as important benchmark systems to investigate ligand binding^70^. In particular, L99A mutant is a well-studied case^71–73^, in which the mutation creates a small artificial cavity that can accommodate benzene and indole derivatives^74–77^. Recently, we showed that the Martini 3 force field can accurately predict the L99A T4 lysozyme ligand-binding pocket and at least four binding pathways^38^. In addition, a nearly quantitative agreement of the binding free energy was obtained for nine different systems including different ligands and the double mutant L99A/M102Q. Given the high similarity of apo and holo states of mutants of T4 lysozyme, which presents a ΔRMSD of 0.2 Å, the system was modeled using the EN approach^38^. However, recent atomistic studies using τ-Random Acceleration MD simulations indicated that maybe such a rigid CG approach was not fully adequate^78^. In particular, it seems that ligand dissociations can involve intermediate metastable protein conformations, which can possibly impact dissociation pathways and rates^78^. Our main hypothesis for this discrepancy was the limited flexibility of the EN approach, which possibly suppressed the small and local conformational changes necessary to open the binding pathways in the intermediate metastable states.

In order to verify this idea, we repeated the Martini 3 MD simulations involving benzene binding to L99A T4 lysozyme using our new enhanced GōMartini approach. The main results are presented in Figure 4. A total sampling of 0.9 ms per system was used here, with the GōMartini model calibrated to show an overall flexibility similar to the EN model. Distribution of the average protein backbone RMSF indicates that the GōMartini model was even slightly less flexible (see Figure 4A) than the EN model. However, comparing the RMSF per residue (Figure 4B) shows a slightly different pattern of flexibility, with the GōMartini model showing more rigid helical regions, but a slightly more flexible region around the L99A T4 lysozyme benzene pocket (C-terminal domain on the bottom of the structures displayed in Figures 4B and 4D). This increased flexibility in the pocket seems to indeed have an impact on ligand binding, with clear local minima being observed in the PMF profile obtained with GōMartini (Figure 4C). These are not observed with the EN model. It is worth mentioning that the binding free energy of the global minimum is almost identical between both models. The new local minima observed with the GōMartini model are located at distances of 0.5 and 0.9 nm from the main pocket (located at ∼0.2 nm in the PMF). Free energy estimates based on ligand densities indicate that the local minimum around 0.5 nm is located in a pre-pocket near the dissociation pathways between helices CD and DG, which also seems to be the most populated metastable intermediate for benzene observed in atomistic τ-Random Acceleration MD simulations^78^. This result strongly suggests that the GōMartini approach can better capture subtle conformational fluctuations of the protein that are involved in fit-induced binding mechanisms.

**Figure 4.**
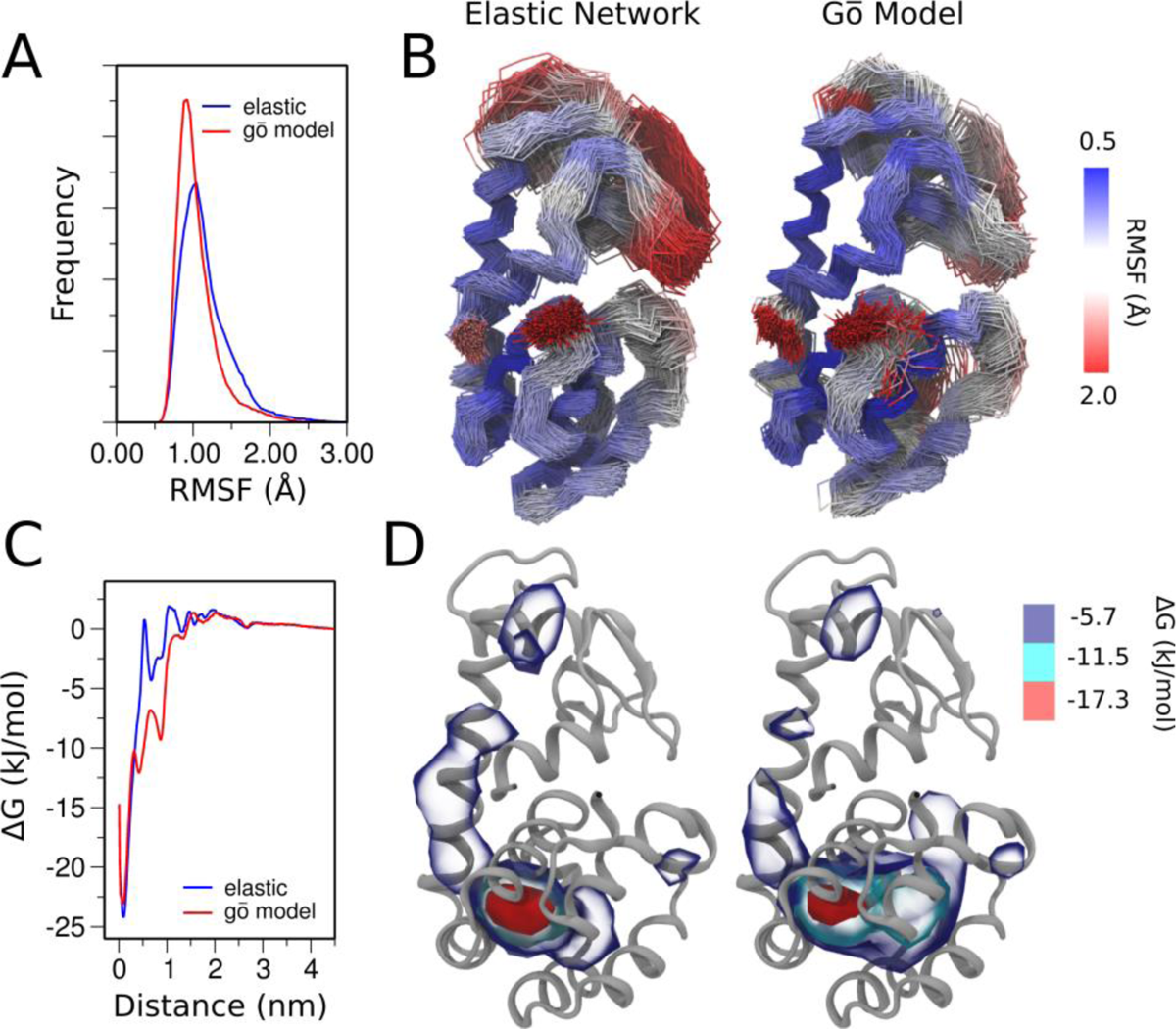
Unbiased simulations of ligand binding to L99A T4 lysozyme with elastic network and GōMartini models. (A) Distribution of the average protein backbone RMSF for the EN (blue) and GōMartini (red) models. (B) Average RMSF per residue of the protein backbone bead (BB) for simulations performed with EN (left panel) and GōMartini (right panel) models. (C) Radial ligand-receptor PMFs obtained with benzene using EN (blue) and GōMartini (red) models. (D) Benzene density around L99A T4 lysozyme obtained from averaging 0.9 ms of CG simulations for EN (left panel) and GōMartini (right panel) models. The blue, cyan and red isosurfaces can be translated to the free energy values shown at the color map.

### SOD1: allosteric pathway detection

Copper-zinc superoxide dismutase (SOD1) is a critical enzyme responsible for catalyzing the conversion of superoxide anions into hydrogen peroxide and molecular oxygen^79,80^. SOD1 has gained significant attention due to its connection with amyotrophic lateral sclerosis (ALS), a neurodegenerative disorder^81^. Over 100 different mutations in the SOD1 gene have been identified as causes for familial variants of the disease^82^, but the precise molecular mechanisms underlying their pathogenicity remain a subject of active debate within the scientific community. It has been proposed that protein aggregation^80^, aberrant pro-oxidant catalysis^83^ and metal dyshomeostasis^84^ may be involved in the pathogenic mechanism. The proposed pathogenic processes listed have in common that they have been linked to loss of Zn(II) ions from the holoprotein^85–91^. Surprisingly, only a subset of ALS-linked SOD1 mutations occur close to the metal site^92^, raising questions about the molecular mechanisms involved in Zn(II) loss.

In its active dimeric form, each SOD1 monomer contains a Cu(I)/(II) ion, critical for catalytic function, and a Zn(II) ion, primarily serving a structural role. Close to these metal ions and the active site is the electrostatic loop (EL), which is known to be destabilized in several ALS-linked mutants^79,93^. We have recently shown that a combination of the virtual site GōMartini approach and the Martini 3 model can provide insights into how subtle structural perturbations in SOD1, induced by mutations such as G93A, located 4 nm away from the catalytic site, might occur. These perturbations could increase the likelihood of the EL detaching from its native position and exposing the metal sites to water. Through extensive 480 μs CG MD simulations for both wild-type and G93A mutant SOD1, an allosteric pathway was identified explaining how the distant G93A mutation affects the EL^41^. Here, we revisit this system to investigate whether similar results can be obtained using simpler EN models. Figures 5A and 5B reveal that overall flexibility trends in the GōMartini and EN models are comparable for wild-type SOD1. However, the GōMartini model exhibits reduced flexibility in the β-barrel core compared to the EN model, while the EL region displays the opposite trend. Strikingly, flexibility comparisons between wild-type and G93A mutant (Figures 5C and 5D) demonstrate that the GōMartini model presents a more complex profile of the RMSF difference with increased stabilization around the mutation site and higher flexibility in multiple loops, including the EL. To elucidate the allosteric pathway through which these changes happen, we found that there were differences in residue-residue distance distributions connecting the mutation site and the EL when G93A mutant and wild-type are compared when using the GōMartini model (Figure 5F). In contrast, our EN model failed to identify any differences connecting the mutation site and the EL. These results highlight the superior capability of Gō models in capturing subtle structural dynamic changes. Moreover, they suggest that the GōMartini approach has a promising potential to study long-range alterations in dynamics induced by single point mutations, even for ones introducing subtle molecular modifications such as the addition of a single methyl group, as exemplified by the G93A mutation of SOD1.

**Figure 5.**
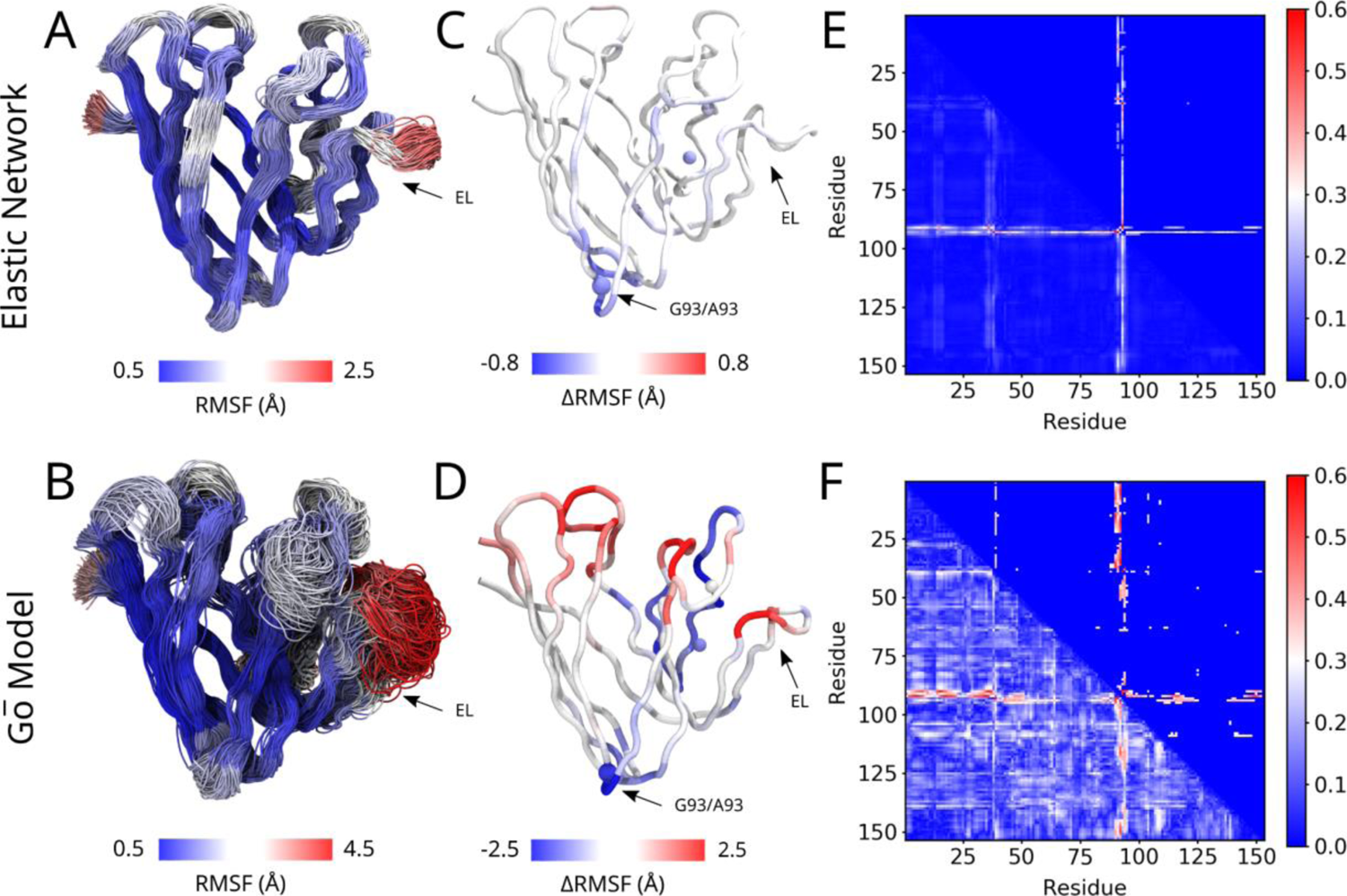
Comparison of the effect of elastic network and Gō models in the allosteric pathways of SOD1. (A and B) Flexibility of the protein backbone of the WT with the elastic network (A) and GōMartini (B) models. Snapshots were taken every 1 μs. The color scale represents the average backbone RMSF per residue. (C and D) Change in RMSF between the WT and G93A with the elastic network (C) and GōMartini (D) models. Blue indicates rigidification in G93A. Red indicates increased flexibility. (E and F) Matrix representation of the integrated absolute difference in the distance distributions between all backbone beads. Results are presented for EN (E) and GōMartini (F) models. The bottom left triangle represents the full data set. In the top right triangle, only values of >0.3 are depicted; all other values are colored blue.

### RBD:H11-H4 complex: Probing the mechanical stability of an antigen:nanobody

The mechanical stability of proteins in enveloped viruses is of great relevance for virus-cell interaction^94^ as emphasized for instance in studies on the SARS-CoV-2 spike (S) protein^95^. SMFS experiments played a key role in unveiling this relevance and in enhancing our understanding of the molecular evolution of the SARS-CoV-2 variants^96^. The key region of the S protein that is associated with cellular recognition is the so-called receptor binding domain (RBD) and in particular this protein domain has presented key mutations in each of the variants of concern that enhanced binding affinity of the entire S protein to the cellular receptor.

Here, we employed the GōMartini approach for probing the interaction of an RBD-nanobody at lower pulling speed than typically accessible by AA MD simulations and furthermore, we avoided to apply position restraints on the RBD as they do not correspond to typical AFM-SMFS protocols. The GōMartini steered MD (SMD) simulations were conducted under similar conditions (without restraints), and we only fixed the position of one residue in the RBD whereas the pulling residue was part of the nanobody. We calibrated the strength of the LJ potential (*ε*LJ) in the GōMartini model following the AA SMD studies by Nguyen and Li^97^. The GōMartini model was applied for both proteins as well as to define the protein complex interface. Note that in the AA reference study position restraint potentials along all BB atoms in the RBD were applied and a very high pulling speed was employed compared to the SMFS experiments. In this regard the GōMartini SMD simulations reproduced quite well the average value of the rupture force, Fmax, using the same pulling speed. Restraining the positions of certain groups of atoms is not equivalent to AFM-SMFS experiments and it is only a convenient way to avoid the protein unfolding in AA MD simulations. Thus, the use of a larger MD simulation box is recommended to capture the full dissociation process. Such simulations have a high cost in AA MD and thus large protein complexes undergoing conformational changes still suffer from limited sampling in SMD. We removed all artificial position restraints and performed the same study using the GōMartini model. The next step was to assess the impact of these artificial position restraints on the nanomechanics of protein complexes. Our average rupture force is about ∼300 pN below the value reported by Nguyen and Li (Fmax ∼ 925 pN) (see Table 1 and Figure 6A).

**Figure 6.**
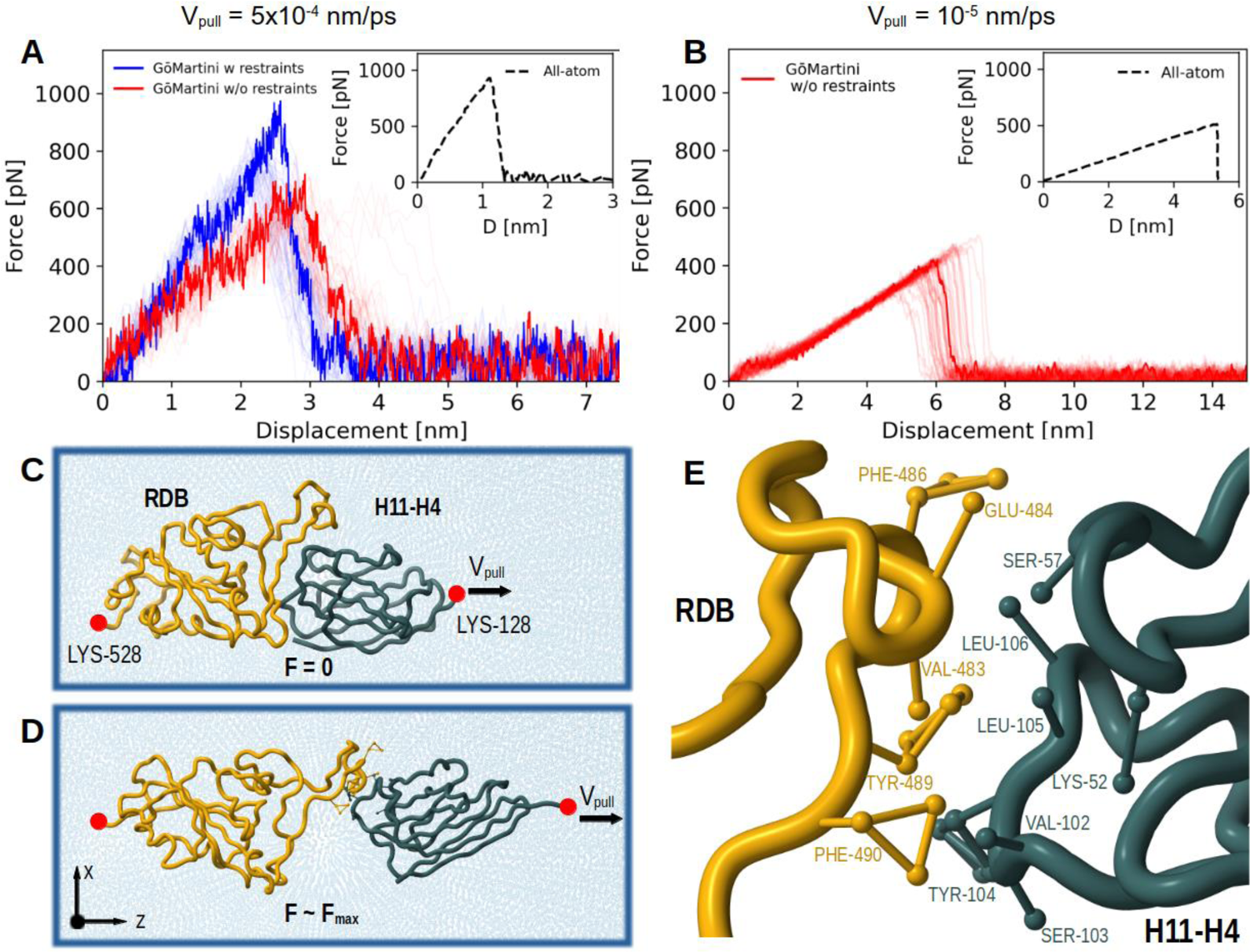
Nanomechanics of the RBD:H11-H4 complex studied by GōMartini simulations at different pulling speeds. (A) Force-displacement profiles for RBD:H11-H4 complex at vpull = 5×10^-4^ nm/ps in SMD simulations using the GōMartini model, with and without position restraints of the BB beads of the RBD. The pulling SMD spring constant was set to 600 kJ/(mol·nm^2^). (B) Same as in (A), but the dissociation of the complex is carried out at vpull = 10^-5^ nm/ps and the pulling SMD spring constant set to 60 kJ/(mol·nm^2^). The inset in A and B shows the reference AA SMD data, note that the y-axis shows the distance (D) between center of mass of groups pulled in AA SMD protocol whereas in GōMartini study the displacement is associated with increase of z value along the pulling direction. (C) Structure of the RBD:H11-H4 complex placed in a box of CG Martini water represented as blue beads in the initial bound state with F = 0 pN. The fixed LYS-528 residue in the RBD and the LYS-128 residue in H11-H4 used for pulling are highlighted by red beads. (D) Structure of the complex at Fmax ∼ 434 pN. (E) Magnified view of the last protein segments in contact before the full dissociation of the protein complex at d ∼ 6 nm. The structures in (C)-(E) are taken from a replica simulated with vpull = 10^-5^ nm/ps.

**Table 1.** Nanomechanical characterization of the RBD:H11-H4 complexes with and without position restraints at different pulling speed (vpull) in SMD. The number of replicas is given next to Fmax values.

| SMD ($v_{\text{pull}}=5 \times 10^{-4}$ nm/ps, $k_b = 600$ kJ/mol·nm<sup>2</sup>) | $F_{\max}$ (pN) |
| --- | --- |
| CHARMM36<sup>97</sup> w/ restraints + | 926±15 (n= 5) |
| GōMartini w/ restraints | 946±75 (n= 50) |
| w/o restraints | 664±45 (n= 50) |
| SMD ($v_{\text{pull}}=10^{-5}$ nm/ps, $k_b = 60$ kJ/mol·nm<sup>2</sup>) | |
| CHARMM36<sup>98</sup> w/ restraints | 508±136 (n=8) |
| GōMartini w/o restraints | 413±43 (n=50) |

In fact, we showed how restraints have a negative impact on the mechanical stability of the protein complex, as they will overstabilize the protein complex which then will not give comparable results with SMFS experiments. In a second study by Golcuk et al. ^98^, position restraints were applied on a smaller number of non-hydrogen atoms located at the RBD:H11-H4 interface. This resulted in the rupture force being similar to the one obtained from our unrestrained GōMartini simulations at a lower pulling speed (see Figure 6B), which is still about four orders of magnitude larger than the typical pulling speed in SMFS experiments (∼10^-9^ nm/ps)^96^. Note that in the AA SMD studies, a handful of replicas were performed whereas at the CG Martini level of resolution a large number of replicas can be run at the same computational effort as the few AA SMD replicas. The nanomechanical characterization pulls at constant speed one side of the complex (i.e. nanobody) while the RBD remains anchored in space and this process perturbs the bound conformation of the H11-H4 nanobody starting from the pulling direction involving residues around LYS-128 (see Figure 6C-D). The analysis of the protein chain at Fmax revealed the stretching of the receptor binding module (residues 424-495), which is the region that is mostly in contact with the ACE2 receptor, such that part of the RBD is perturbed by the nanobody before dissociation. We identified several hydrophobic interactions as the most relevant ones for the buildup of Fmax: VAL-483/SER-57, GLU-484/LYS-52, PHE-486/LEU-106, TYR-489/SER-103, TYR-489/TYR-104, and PHE-490/VAL-102 (see Figure 6E). An additional nanomechanical study was performed with the dockerin:cohesin protein complex system, with the results displayed at the Supporting Results C2. Our nanomechanical profiles captured the two most prominent dissociation pathways observed in by previous all-atom SMD simulation^99^.

## 6. PERSPECTIVES: HOW TO IMPROVE GŌMARTINI AND THE PROTEIN MODEL

### Improving contact maps and strength of interactions

The combination of a Gō-like network with the Martini CG force field can be effectively employed to capture conformational changes. However, the choice of parameters to build the network is not obvious. To address this question, we explored the possibility of improving the key parameters of the GōMartini model: the strength of the interactions (*ε*LJ) and the contact map. A convenient possibility is to fine-tune these parameters based on AA MD simulations instead of a single experimental structure. Nonetheless, it is worth noting that the initial GōMartini model can still exhibit a bias due to its starting configuration, thereby directing the simulations towards the native conformation. A common issue encountered in this context pertains to the definition of unnecessary Gō bonds within loop regions, mainly attributed to the tightly packed nature of these regions in the crystallographic and cryo-EM structures. Consequently, the native contacts may underestimate the flexibility of loop regions. Employing dynamical contact analysis of the AA MD simulations could distinguish between stable and transient contacts within the protein structure on the timescale of the AA simulation.

As a first exploration of how GōMartini parameters could be refined considering a dynamic contact analysis, several benchmark studies, including soluble (as per previous work by Poma et al.^34,36,37^) and transmembrane proteins ranging from 76 to 4160 residues were explored here. We initially focused on optimizing the effective depth of the LJ potential (*ε*LJ) while preserving all Gō potentials, aiming to bring the standard GōMartini model in closer agreement with the protein dynamics observed in AA simulations, particularly in terms of RMSF. Notably, the optimal *ε*LJ value exhibited significant variation across different systems, highlighting the importance of tailoring *ε*LJ values to individual protein models rather than employing a uniform value across all systems.

As shown in Figure 7, we computed the RMSF for the Cα atoms and BB beads in the AA and CG models, respectively, for three benchmark proteins: titin I-band (1TIT), glycoside hydrolase (3W0K), and the transmembrane domain of Ist2. Upon comparison of the original GōMartini model (blue lines) with the CHARMM36 AA reference (black), it becomes evident that although using specific *ε*LJ values calibrated for each protein significantly improved the original GōMartini models, they still fail to capture the dynamics of several loop regions observed in AA simulations. Indeed, RMSF analysis indicates that these regions remain relatively rigid in standard GōMartini models compared to AA simulations.

**Figure 7.**
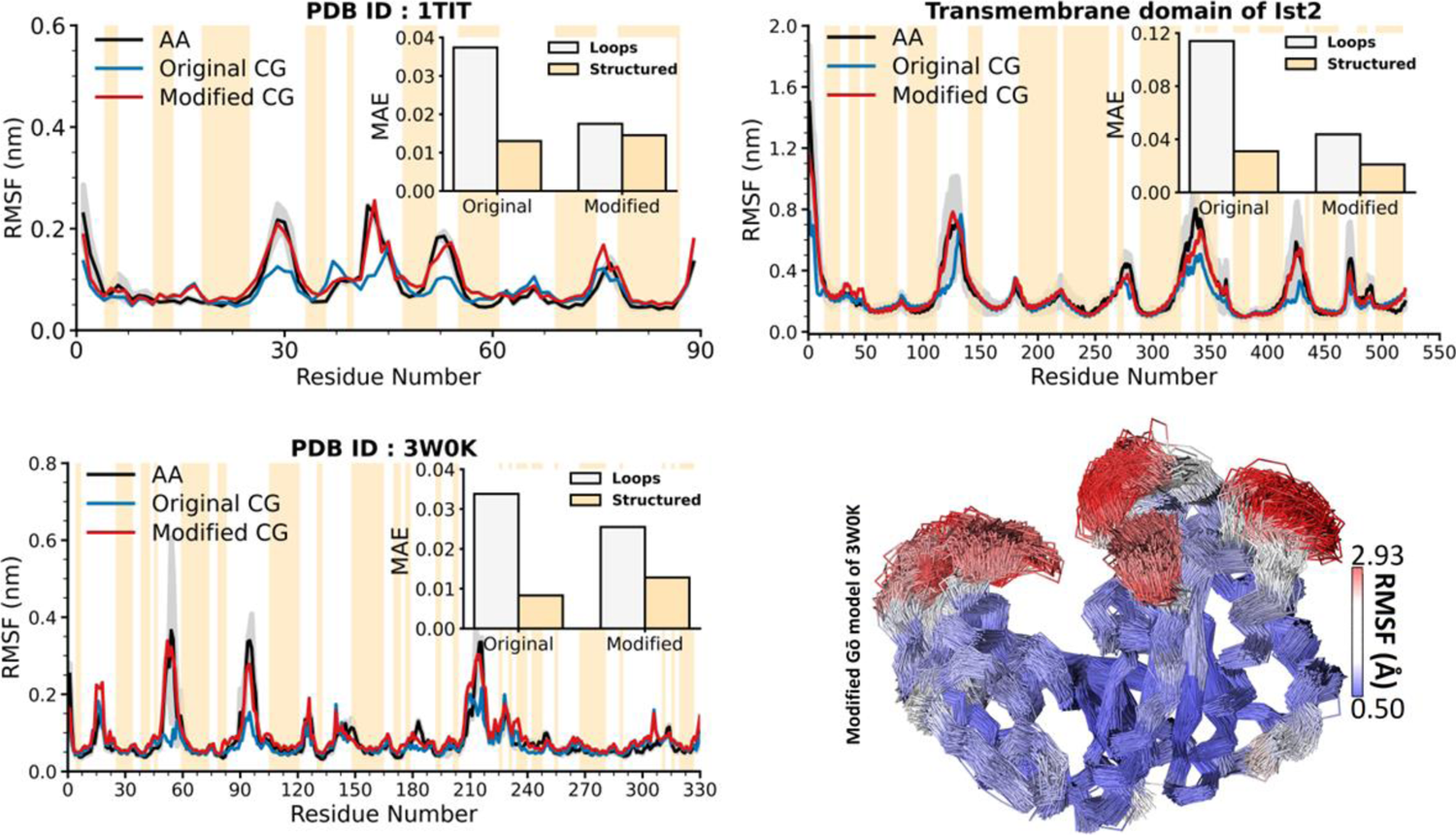
Improving GōMartini to match AA models. RMSF comparison between original GōMartini (with *ε*LJ optimized), modified GōMartini (with removal of Gō interactions in loops) and AA simulations for three proteins: titin I-band (1TIT), glycoside hydrolase (3W0K), and the transmembrane domain of Ist2. The mean absolute error (MAE) for loops and structured regions between GōMartini and AA simulations is highlighted in the insets. The bottom-right panel presents the flexibility of the protein backbone beads during simulations using the modified GōMartini model for glycoside hydrolase (3W0K).

To overcome this challenge and improve the accuracy of the GōMartini models, we further optimized them by checking the contact frequencies around each residue throughout an AA MD simulation (see the Supporting Methods B5 for more details). The results of this optimization are summarized in Table S2. Figure 7 shows a good agreement between the modified GōMartini model (red lines) and the CHARMM36 AA reference (black), which demonstrates that the modified GōMartini model accurately captures characteristic fluctuations across most residues and is flexible enough to mimic the flexibility observed in AA MD simulations. It is important to highlight that, in the case of ubiquitin, cohesin, and aquaporin (see Figure S5), the method did not substantially improve the flexibility of the protein. Overall, it appears that investigations of systems featuring substantial conformational transitions require fine tuning of Gō networks, shown here, and the possibility of flexible secondary structure, not included in current models.

### Go virtual-sites for adding water-bias in IDPs and biomolecular condensate

While the Martini 3 protein model has already greatly improved upon the previous iteration, two years after release, room for improvement has been identified regarding some specific aspects of the model. Studies have reported that Martini 3 underestimates the radius of gyration (Rg) of IDPs and multidomain proteins in solution when compared to experimental SAXS data^44,100^. Simultaneously, it has also been shown recently that the behavior of transmembrane domains might be unstable in Martini 3, specifically in the case of transmembrane alpha helical peptide insertion^45,101,102^. For both cases, scaling of protein-water interactions, which was a common mitigation strategy employed in Martini 2, was suggested to resolve the issues. Thomasen et al.^100^ found that increasing protein-water interactions by 10 % results in improved agreement with SAXS data for IDPs and multidomain proteins, while Cabezudo et al.^101^ found that reducing protein-water interactions by 10 % resulted in the correct insertion of transmembrane peptides. However, scaling interactions has the major downside of impacting all pair interactions that were altered, and not just the ones responsible for the unintended model behavior — i.e. scaling the P2-W pair interaction, impacts not only protein BB-water interactions but also the interactions involving any other molecule containing P2 beads. This causes major transferability issues for the model, and as such, should be avoided if possible.

Although the virtual interaction sites built on top of BB beads are typically used only to define interactions to other sites for tertiary structure preservation, they additionally offer the possibility to effectively modify interactions between BB beads and other Martini beads in a site-specific manner. For example, by defining an interaction between a BB virtual site and water beads, it is possible to effectively increase the strength of the interaction between protein backbones and water. The strength of the resulting non-bonded interaction will be the sum of the P2-W and virtual site-W interactions. As the interaction is defined only between the virtual site and water beads, the increase in the strength of this interaction is restricted to P2 beads in the protein backbone only, and no other molecules are affected. Further, this approach is sufficiently versatile that it can be applied in only specific residues of proteins, such as transmembrane or disordered domains.

To showcase this, we tested the Rg of select IDPs using the new GōMartini implementation. We used the set of IDPs from Thomasen et al.^100^ to validate our approach against an existing one. The IDPs were coarse-grained and virtual Gō sites included on the BB particles. An additional LJ interaction between the virtual Gō sites and water beads was added, with ε = 0.5 kJ/mol, which when summed to the already existing BB-water interaction roughly corresponds to a 10 % increase of the interaction. Although this value seems the same as proposed by Thomasen et al.^44,100^, it is only applied to backbone-water interactions while previous approaches applied the changes to the whole protein.

To further improve the model, we also developed a refined set of bonded parameters for backbones and side chains using AA simulations, as these are not implemented for coiled structures in *Martinize2* (Figure S7). The addition of either component individually goes some way to improving the radius of gyration of the target set of IDPs, reducing the mean absolute error with respect to the experimental reference across the set from 1.35 nm to 1.25 nm in the case of the additional bonded parameters, and to 0.36 nm with the addition of the Gō site dedicated to water interactions (Figure S8). However, as we show in Figures 8A-B, the combination of these extra parameters together further improved the Rg of the benchmark set with respect to the experimentally measured values, with a final mean absolute error of 0.28 nm.

**Figure 8.**
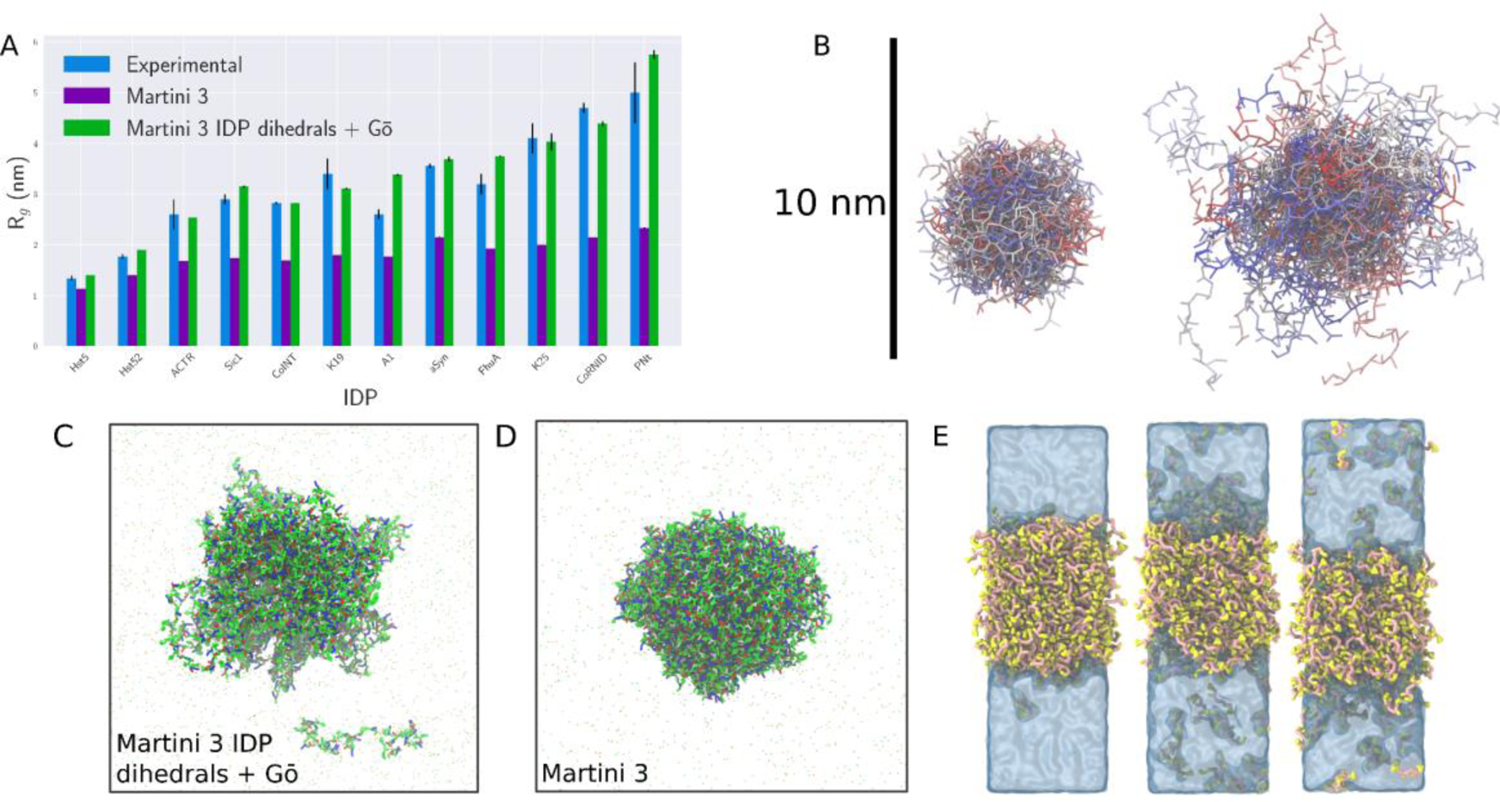
Improving IDP global dimensions and condensation using Martini 3 with GōMartini model-based interaction rescaling. (A) Radii of gyration of the IDP benchmark set of Thomasen et al. Results are compared between the experimental value (blue) to both the native Martini model (purple) and optimized Martini IDP + Gō model with additional bonded parameters (green). (B) Illustrations of the increase in ensemble dimensions of ACTR comparing (left) native Martini 3 and (right) the final model for IDPs with additional bonded and non-bonded potentials. (C,D) Illustrative snapshots of a condensate (with improved IDP parameters) and an aggregate (with default Martini 3 parameters) of an artificial IDP (WT20) known to phase separate. (E) Snapshots of FLssLF peptide systems with varying increases in the strength of the BB-water interactions. Left; native Martini (0% increase); middle: 6% increase; right: 8% increase of protein BB-water interactions.

To additionally validate the use of virtual sites in improving the behavior of Martini IDPs, we carried out simulations of a known phase separating IDP. Recent work from Dzuricky et al. designed an artificial IDP that would demonstrate liquid-liquid phase separation based on an octapeptide repeating unit^103^. In Figure 8C, we show a phase separated system of this protein after 5 µs, where the optimized IDP model above has been used for the WT20 construct (ie. 20 octapeptide repeats using the primary so-called ‘wild-type’ sequence). In contrast, Figure 8D shows that without the optimization of the IDP described above, the phase separation of the model is visibly different, being more compact. This system in fact does not form a liquid-like condensate, but a solid-like aggregate, as evidenced by the analysis of the incoherent scattering curves in Figure S9.

As a further example of how increasing protein BB-water interactions can aid recapturing experimental behavior, we simulated a system of two FL dipeptides linked by a disulfide moiety. This system was shown to undergo liquid-liquid phase separation in the recent work of Abbas et al^104^. Figure 8E shows that native Martini 3 could capture the condensate formation, with a coexisting dense and a dilute phase. However, the resulting condensate was too dry (∼10% water weight content) compared with the experimental data (∼62%). To alleviate this problem, again we introduced virtual Gō sites on the BB beads, carrying an additional interaction with the water beads. As shown in Figure 8E, increasing the BB-water interactions increases the water content of the condensate without affecting the phase separation. With an 8% increase in the strength of this interaction (corresponding to ε = 0.3464 kJ/mol), the water weight content already is above 50%, much closer to the experimental findings. Overall, these results demonstrate that beyond a universal rescaling, the strength of the BB-water interaction can be fine-tuned to better reproduce properties of biomolecular condensates.

### Gō virtual sites for adding water-bias in TM helices and beta-sheet peptides

As mentioned in the previous section, there have been reports of issues surrounding the transmembrane insertion of some helical peptides using Martini 3^45,101,102,105^. The solution previously proposed to overcome these issues was again to apply a rescaling of peptide-water interactions, similar to what has been done for IDPs. Here, we have also tested our GōMartini implementation, changing only the BB-water interactions. Four WALP α-helices – 16, 19, 23, and 27 residues in length, termed WALP16, WALP19, WALP23, and WALP27 – were coarse-grained using our GōMartini implementation and simulated embedded in a dimirystoylphosphatidylcholine (DMPC) membrane, as done by Spinti et al. ^45^. To facilitate the observation of WALP ejection from the membrane, a temperature of 310 K was used, instead of the 300 K used by Spinti et al. ^45^. A second set of these systems was run where an additional LJ interaction between the virtual Gō sites in helical residues and water beads were included, with ε = −1.0 kJ/mol, so effectively reducing the interactions with water to improve peptide insertion. The reduced LJ interaction substantially stabilized the transmembrane conformation of the four WALPs and reduced the TM peptide ejection in comparison to the control simulations (Figures 9A-B, Figure S10).

**Figure 9:**
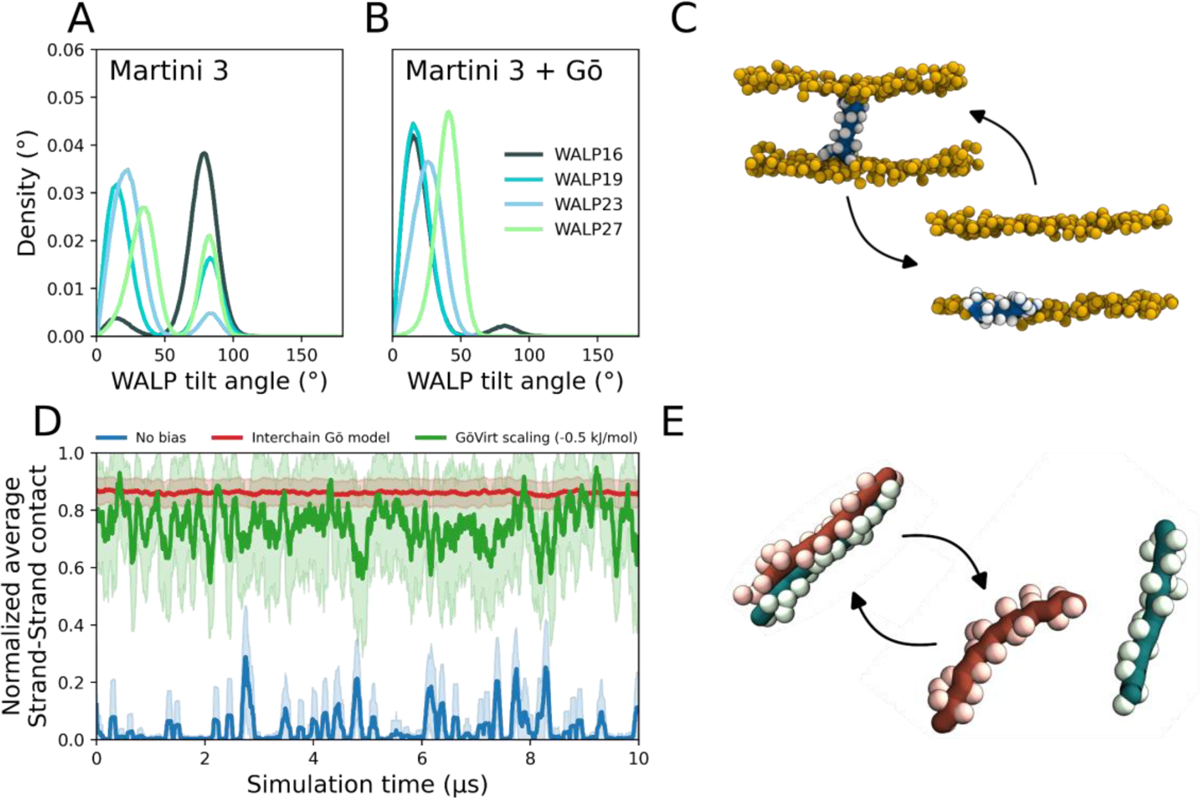
Improving transmembrane peptide insertion and beta-sheet aggregation. Tilt angle distributions from simulations of WALP peptides inserted in DMPC membranes using the GōMartini model with either (A) no additional LJ interaction, or (B) an additional LJ interaction between the virtual Gō sites and water beads of ε = −1.0 kJ/mol; tilt angle states close to 0° correspond to TM configurations, whereas those close to 90° correspond to peripherally membrane-adsorbed ones. (C) Representative WALP16 configurations, both fully inserted in its preferred transmembrane configuration and in its peripherally membrane-adsorbed state. WALP16 backbone shown in blue, with side chains in white. Membrane phosphate beads are represented in orange. (D) Normalized average contacts between two RAD16-I peptide beta strands. Solid lines show the running averages of 500 frames, while the shaded area shows the running standard deviation. Simulations were run with a GōMartini model applied between the two chains (red), with an additional LJ interaction between the virtual Gō sites and water beads with ε = −0.5 kJ/mol (green), and without any structural or interaction bias (blue). (E) Representative RAD16-I strand configurations, both aggregated and dissociated. Each backbone chain is colored in either brown or green, with the side chains colored in a lighter shade of the same color.

Given the need to rescale BB-water interactions with both helical and coil protein segments, we aimed to assess whether beta strand segments could also benefit from the rescaling of their BB-W interactions. To do so, we tested how the aggregation of RAD16-I is currently performing with Martini 3. RAD16-I is a synthetic amphipathic peptide that adopts a beta-strand structure in solution. It has been shown that RAD16-I associates to form a very stable beta-sheet in solution, even culminating in the formation of nanofibers with increasing peptide concentration.^106,107^ We assembled a simple system containing 2 copies of RAD16-I and followed their aggregation into a 2-strand beta sheet over the course of the simulation (Figure 9, D-E). As expected, with a GōMartini applied between the two chains we can accurately reproduce the 2-strand beta sheet assembly. However, in the absence of any bias, we do not observe stable sheet assembly. To mitigate this, we applied an additional LJ interaction between the virtual Gō sites in strand residues and water beads with ε = −0.5 kJ/mol. This reduction of BB-W interactions was sufficient to obtain sheet assembly similar to the one observed with the interchain Gō model.

## 7. DISCUSSION

Despite attempts to explore multiple-state conformations states^108^ or to enhance the accuracy of Martini protein models through approaches such as polarizable^26,109^, titratable^110,111^, and even possibly foldable^109^ versions, the combination of standard Martini with a bias, such as an elastic network or a Gō model, remains the most attractive and useful option due to its computational performance and compatibility with large libraries of Martini models.

The virtual-site implementation of Gō models with Martini 3 was initially introduced as a proof of concept in the works involving SOD1^41^ and light-harvesting complex II^35^, being officially adopted as the Gō approach tested during Martini 3 development. Since the release of Martini 3, the model has been recommended and included in our tutorials^112^ and thus been extensively tested by the modeling community. However, the key features of the approach and the underlying Martini 3 protein model had not been presented until now. The main goal of this work was to finally detail all the advantages of the current implementation in relation to elastic networks and the previous GōMartini implementation. Concerning the advantages in relation to elastic networks, we clearly show here how the improved conformational flexibility — stemming from the use of asymmetric potentials with finite dissociation energies — can be used to study long-range allosteric changes in proteins, protein-small molecule binding, and protein-membrane binding. These kinds of applications are still new within the Martini community and we foresee more studies in the future showing the benefits of the more accurate protein flexibility introduced by GōMartini. However, it is important to highlight that this approach is currently not suitable for systems consisting of many copies of the same protein, given that the same Gō bonds which stabilize certain folded states will also wrongly impact protein-protein interactions. Moreover, while different protein aliases can be used for the different copies of the same protein to remedy the impact on protein-protein interactions, the number of protein-protein interface combinations may possibly explode beyond what can be handled by this approach. For instance, a simulation box with 100 copies of the same protein, with each protein with two protein-protein interfaces, each with 10 contacts, would need to have a total of 10*(100!/(100-2)!) = 99,000 interface contacts to be defined. Thus, simulations of crowded membranes^8^ and even the future cell-simulations^113^ may still rely on simple elastic network approaches.

Although nanomechanics studies have been performed before with the previous GōMartini implementation, our results here reinforce its accuracy and show some advantages in reproducing conformational transitions for simulations mimicking AFM profiles. The first GōMartini implementation by Poma et al. captured the unfolding profile of the I27 domain of titin, type I cohesin domain, and ubiquitin with experimental forces equal to 204 pN, 480 pN, 230 pN respectively^34^. Although GōMartini simulations correctly reproduced the expected trends for these systems, the forces were twice as large due to the speed of the SMD simulation (e.g. ∼10^-3^ nm/ps). In this regard, the current virtual site implementation is more convenient as it allows for the full integration of GROMACS^60^ and OpenMM^114^ parallelization and thus, one can use lower pulling speeds with the SMD protocol of ∼10^-5^ nm/ps, which is significantly closer to the pulling speed of SMFS experiments of ∼10^-9^ nm/ps, without compromising computational cost.

In addition to the gain in computational performance and numerical stability in relation to previous versions, the use of virtual sites provides the flexibility to introduce corrections to the backbone-water interactions. In contrast to recently published approaches, we suggest using only water interaction biases in relation to the backbone beads. One of the key reasons is the overall quality of the water/oil partitioning estimates of side chains (Figure S1 and Table S1), which does not show any particular trend of being too hydrophilic or too hydrophobic, with average errors below ∼3 kJ/mol. On the other hand, the hydrophobicity of the protein backbone has always been under debate, as its capability of forming internal hydrogen bonds in secondary structure motifs may affect its partitioning to different environments. This idea was one of the main assumptions of the original Martini 2 implementation^25^, and was recently also incorporated in the SPICA CG model^115,116^. Although some of our results presented here may point out that the Martini 3 model could benefit from the same approach, a broader view of peptides and proteins in different contexts indicates that water biases dependent on the secondary structure are not general. For instance, soluble globular proteins with high helical content, such as lysozyme^31^, may aggregate too much if we consider the correction indicated for transmembrane proteins in this work. In other cases, such as protein/peptide dimers, the solution may be beneficial, although a direct application of an interchain Gō network is more accurate. Indeed, in the case of protein complexes, several studies^36,37^ reported the need to model the complexes with additional interchain Gō bonds at the interface either using the GōMartini approach or alternative Gō models (i.e., OLIVES^117^). The combined representation of structure-based models at the interface of protein complexes led to capture large conformational changes under nanomechanical probing^36,37^, as it is studied by SMFS. Many aspects of the current protein model are being revisited now, including further improvements in side chain self-interaction, improved backbone torsions and side chain rotamers, and more detailed backbone models. These improvements take advantage of the new features of Martini 3, including different bead sizes and labels.

One additional aspect touched by our work is the possibility of refining GōMartini parameters, i.e. the depth of the potentials and contact map. We show that the use of specific potential depths *ε*LJ and improvements in the contact map based on multiple reference structures, obtained for instance from atomistic simulations, can greatly improve the overall flexibility of the models. Automatic refinement of these parameters may be possible via approaches based on particle swarm optimization strategies such as CGCompiler ^118^ and SwarmCG^119^. Similar ideas can be developed considering AI-based approaches instead of atomistic MD references, such as the recent implementation of elastic networks using AlphaFold confidence scores^29^. It is promising to further expand this kind of approach in the future, because it may be the key for the simultaneous representation of multiple conformational states. One recent implementation in this direction involves replacing the single-basin Gō model with a multiple-basin Gō model^120^ or at least a double-well potential. This modification may allow large conformational changes, enabling a more accurate representation of the transitions between stable folded states of proteins.

A last important remark regarding the new virtual site approach is related to our view for future protein model development in the Martini force field: development of Martini protein models and bias approaches, such as GōMartini and elastic network models, should be decoupled. In previous iterations, these two aspects were interconnected in a way that biases in secondary and tertiary structure were fully integrated in the model, even affecting bead types and mapping^25,28^. Such integration blocked further development, as any attempts to change the model would need to involve developing both the core model and the bias. For instance, improving protein flexibility to allow secondary structure changes in Martini 2 would depend on dramatically changing fundamental aspects of the model as bead types depend on secondary structure, and elastic networks were of paramount importance for beta-sheet stability. Therefore, we advocate for a complete decoupling of the two developmental pathways. While the protein models should follow the typical building-block rules and validation of Martini models, the structural biases should always come as an additional experimental/theoretical potential applied on top of the model, used to bias the simulated ensembles. This approach guarantees that the Martini protein model can independently evolve, with further improvements hopefully resulting in the use of less/weaker biases. Although the ultimate aspiration remains the creation of a bias-free Martini protein model, we recognize the enduring importance of approaches such as GōMartini as fundamental tools for accurately modeling proteins within the Martini universe.

## ASSOCIATED CONTENT

**Supporting Information**. A listing of the contents of each file supplied as Supporting Information should be included.

## AUTHOR INFORMATION

### Author Contributions

P.C.T.S. and S.T. conceived and initiated the project, with later contributions in the design of the project from S.J.M. and A.B.P.. L.B.A., F.G., and S.T. wrote the codes. L.B. and P.C.T.S. reviewed and the detailed the Martini 3 protein model, including compilation of the partitioning free energies and SASA of the side-chain analogues. S.T. contributed to the studies involving the PH domain of PLCδ1. P.C.T.S. contributed to the studies involving the T4-lysozyme and SOD1, with additional contributions of R.M.A. and S.T. for the SOD1 work. R.A.M., L.F.C.V., and A.B.P. contributed to the nanomechanic studies. H.R., L.M., and P.C.T.S. contributed to the flexibility improvement benchmarks via refinements in the contact map and strength of interactions. C.B., L.W., P.P., and P.C.T.S contributed to the IDP and biomolecular condensate studies. L.B.A., A.C.B.A., S.W., M.N.M., and P.C.T.S. contributed to the transmembrane peptide studies. P.C.T.S. and S.T. wrote the manuscript with contributions from all the authors. All authors discussed the results, revised the manuscript, and approved the final version of the manuscript. S.J.M., S.W., P.C.T.S, and A.B.P. provided most of the financial and computational resources of the project, with contributions from L.M. and S.T..

### Notes

The authors declare no competing financial interest.

## ACKNOWLEDGMENT

This work was granted access to the HPC resources of IDRIS and TGCC under the allocations 2022-A0120713456 and 2023-A0140713456 made by GENCI. We also acknowledge the support of the Centre Blaise Pascal’s IT test platform at ENS de Lyon (Lyon, France) for the computer facilities. The platform operates the SIDUS solution developed by Emmanuel Quemener^121^. We thank the Center for Information Technology of the University of Groningen for providing access to the Peregrine high-performance computing cluster. We also acknowledge the National Computing Facilities Foundation of The Netherlands Organization for Scientific Research (NWO) for providing computing time. L.B.A and P.C.T.S would like to thank the support of the French National Center for Scientific Research (CNRS) and the funding from research collaboration agreements with PharmCADD. S.J.M. received funding from the European Research Council (ERC) through an ERC Advanced grant “COMP-MICR-CROW-MEM”. S.T. acknowledges the support from the European Commission via a Marie Skłodowska-Curie Actions individual fellowship (MicroMod-PSII, grant agreement 748895), the Center for Multiscale Modelling in Life Sciences (CMMS), the Alfons und Gertrud Kassel Foundation, and the Dr. Rolf M. Schwiete Foundation. L.M. acknowledges funding by the Institut National de la Santé et de la Recherche Médicale (INSERM). M.N.M. acknowledges Fundação para a Ciência e a Tecnologia for fellowship CEECIND/04124/2017/CP1428/CT0008. A.B.P and R.A.M. acknowledge Marek Cieplak for sharing the source code of the OV+rCSU contact map in Fortran. P.P. acknowledges FAPESP support (grant 2019/26557-8). A.B.P. acknowledges financial support from the National Science Center, Poland, under grant 2022/45/B/NZ1/02519 and gratefully acknowledge Polish high-performance computing infrastructure PLGrid (HPC Centers: ACK Cyfronet AGH) for providing computer facilities and support within computational grant no. PLG/2023/016519. R.M.-A. thanks ANID-Chile for financial support under FONDECYT N. 1200200.

