## Supporting Information for "GōMartini 3: From large conformational changes in proteins to environmental bias corrections"

* Authors for correspondence:

| | |
|---|---|
| Paulo C. T. Souza. | |
| Adolfo B. Poma | |
| Siewert J. Marrink | |
| Sebastian Thallmair | |

**SUMMARY**

(A) SUPPORTING NOTES 3

(A1) Martini 3 protein model 3

(A2) Format of the contact map file 5

(B) SUPPORTING METHODS 6

(B1) PH domain of phospholipase C δ1 6

(B2) T4 lysozyme L99A mutant 6

(B3) Wild-type and G93A mutant of Copper, Zinc Superoxide Dismutase 1 7

(B4) Nanomechanics of protein complexes by GōMartini model 7

(B5) Protein flexibility benchmarks 9

(B6) Intrinsically disordered proteins 12

(B7) Biomolecular condensates 13

(B8) Transmembrane peptides 14

(B9) RAD16-I peptide 14

(C) SUPPORTING RESULTS 16

(C1) PLCδ1 PH domain: PMFs for strong lipid binding 16

(C2) Assessing the mechanic stability of XMod-Coh:Doc complex from bacteria 18

(C3) Improving contact maps and strength of interactions 20

(C4) IDPs and biomolecular coacervates 21

(C5) TM helices and beta-sheet peptides 25

(D) REFERENCES 26

(A) SUPPORTING NOTES

*(A1) Martini 3 protein model*

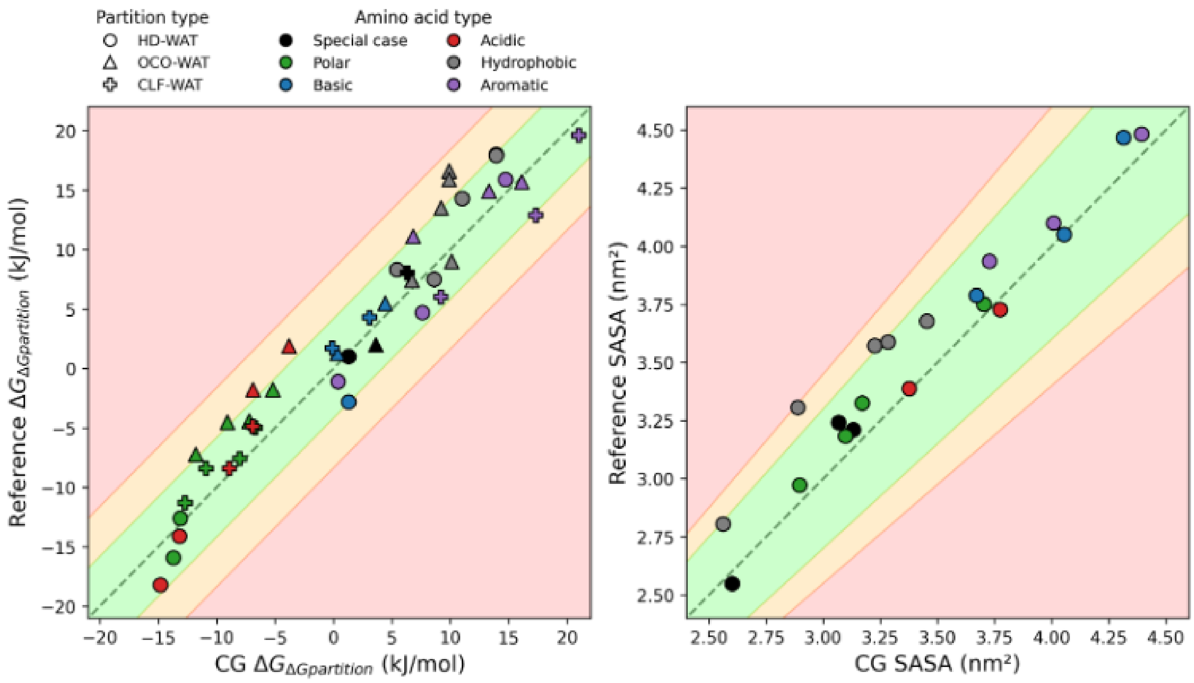


**Figure S1:** Comparison of calculated and reference water–oil partitioning free energies (ΔG) and SASAfor side-chain analogues. For the ΔG, the green, orange and red limits represent 4.184 kJ/mol, 8.368 kJ/mol and over 8.368 kJ/mol deviation from the experimental reference, respectively. For the SASA, the green, orange and red limits represent a 10%, 15% and over 15% deviation from the atomistic simulation reference, respectively.

**Table S1.** SASA and Partitioning of amino acid side chains.

| Category | Amino acid | | *Side chain bead type (charge)* | HD-WAT | OCO-WAT | CLF-WAT | SASA ($nm^2$) |
|---|---|---|---|---|---|---|---|
| Special Cases | GLY (G) | *ref.* | - | - | - | - | 2.548 |
| | | CG | *n/a* | - | - | - | 2.602 |
| | PRO (P) | *ref.* | - | - | - | - | 3.241 |
| | | CG | *SC3* [6] | 11.0 | 9.2 | 10.9 | 3.068 |
| | CYS (C) | *ref.* | *Propane-1-thiol* [1 2 3 4 5] | 7.0 (1.0)[a] | 8.0 (2.0)[a] | 14.0 (8.0)[a] | 3.21 |
| | | CG | *TC6* [6] | 1.3 | 3.6 | 6.3 | 3.132 |
| Polar | SER (S) | *ref.* | *Methanol* [1 2 3 4 5] | -15.9 | -4.4 | -7.6 | 2.972 |
| | | CG | *TP1* [6] | -13.7 | -7.2 | -8.0 | 2.897 |
| | THR (T) | *ref.* | *Ethanol* [1 2 3 4 5] | -12.6 | -1.8 | -5.0 | 3.183 |
| | | CG | *SP1* [6] | -13.1 | -5.2 | -6.7 | 3.098 |
| | ASN (N) | *ref.* | *Acetamide* [1 2 3 4 5] | -26.9 | -7.2 | -11.3 | 3.325 |
| | | CG | *SP5* [6] | -20.4 | -11.8 | -12.7 | 3.171 |
| | GLN (Q) | *ref.* | *Propanamide* [1 2 3 4 5] | -23.0 | -4.5 | -8.4 | 3.75 |
| | | CG | *P5* [6] | -19.2 | -9.1 | -10.9 | 3.704 |
| Basic | ARG (R) | *ref.* | - | - | - | - | 4.467 |
| | | CG (neutral) | - | - | - | - | - |
| | | CG (charged) | *SC3 - SQ3p (+1)* [6] | - | - | - | 4.315 |
| | HIS (H) | *ref.* | *4M-Imidazole* [1 2 4 5] | - | 1.3 | 1.7 | 3.788 |
| | | CG (neutral) | *TC4 - TN6d - TN5a* [6] | - | 0.3 | -0.1 | 3.671 |
| | | CG (charged) | *TC4 - TP1dq (+0.5) - TP1dq (+0.5)* [7] | - | - | - | 3.671 |
| | LYS (K) | *ref.* | *Butylamine* [1 2 4 5] | -2.8 | 5.6 | 4.3 | 4.05 |
| | | CG (neutral) | *SC3 - SN6d* [6] | 1.3 | 4.4 | 3.1 | 4.055 |
| | | CG (charged) | *SC3 - SQ4p (+1)* [6] | - | - | - | 4.055 |
| Acidic | ASP (D) | *ref.* | *Acetic acid* [1 2 3 4 5] | -18.2 | -1.8 | -8.4 | 3.387 |
| | | CG (neutral) | *SP2* [6] | -14.8 | -6.9 | -8.9 | 3.377 |
| | | CG (charged) | *SQ5n (-1)* [6] | - | - | - | 3.377 |
| | GLU (E) | *ref.* | *Propionic acid* [1 2 3 4 5] | -14.1 | 1.9 | -4.9 | 3.727 |
| | | CG (neutral) | *P2* [6] | -13.2 | -3.8 | -6.9 | 3.775 |
| | | CG (charged) | *Q5n (-1)* [6] | - | - | - | 3.775 |
| Hydrophobic Residues – Non-Aromatic | ALA (A) | *ref.* | *Ethane* [1 3 4 5] | 10.5 (7.5)[a] | 10.4 (7.4)[a] | - | 2.805 |
| | | CG | *TC3* [6] | 8.6 | 6.7 | 8.3 | 2.562 |
| | ILE (I) | *ref.* | *Butane* [1 3 4 5] | 18.0 | 16.6 | - | 3.572 |
| | | CG | *SC2* [6] | 13.9 | 9.9 | 11.6 | 3.226 |
| | LEU (L) | *ref.* | *2-methyl-propane* [1 3 4 5] | 17.9 | 15.9 | - | 3.588 |
| | | CG | *SC2* [6] | 13.9 | 9.9 | 11.6 | 3.284 |
| | MET (M) | *ref.* | *dimethyl-sulfide* | 5.3 (8.3)[a] | 6.0 (9.0)[a] | - | 3.677 |
| | | CG | *C6* [6] | 5.4 | 10.1 | 11.4 | 3.454 |
| | VAL (V) | *ref.* | *Propane* [1 3 4 5] | 14.3 | 13.5 | - | 3.306 |
| | | CG | *SC3* [6] | 11.0 | 9.2 | 10.9 | 2.889 |
| Hydrophobic Residues – Aromatic | PHE (F) | *ref.* | *Toluene* [1 2 3 4 5] | 15.9 | 15.7 | 19.6 | 3.935 |
| | | CG | *SC4 - TC5 - TC5* [6] | 14.7 | 16.1 | 21.0 | 3.728 |
| | TRP (W) | *ref.* | *Methyl-indol* [1 2 3 4 5] | 4.7 | 14.9 | 12.9 | 4.482 |
| | | CG | *TC4 - TN6d - TC5 - TC5 - TC5* [6] | 7.6 | 13.3 | 17.3 | 4.394 |
| | TYR (Y) | *ref.* | *para-cresol* [1 2 3 4 5] | -1.1 | 11.1 | 6.0 | 4.099 |

a Corrected value to account for mismatch in carbon chain length. Applied as per https://doi.org/10.1021/acs.jctc.9b00473 .

All the experimental references for water/oil partitioning free energies comes from the following references:[1–5]

*(A2) Format of the contact map file*

The contact file format requires all contacts to be described in a table with 18 columns and the first column to be a capital R. This is the format generated by the web-server http://pomalab.ippt.pan.pl/GoContactMap/[6,7] (which is replacing the previous one[8]: http://info.ifpan.edu.pl/~rcsu/rcsu/) or via the ContactMapGenerator program available at https://github.com/Martini-Force-Field-Initiative/GoMartini. Any information described above the table is discarded. An example generated for ubiquitin (1UBQ) is displayed below.

```
Reading file:    uploads/65ddf463e9bbc/1ubq.pdb
pdb natoms:      602
pdb nresidues:   76
Memory usage:     6.21 MB
UNMAPPED ATOM GLY OXT
Fibonacci grid:  610
ALPHA:           1.24
WATER_RADIUS:    2.80

Residue-Residue Contacts

ID         - atom identification
I1,I2      - serial residue id
AA         - 3-letter code of aminoacid
C          - chain
I(PDB)     - residue number in PDB file
DCA        - distance between CA
CMs        - OV , CSU , oCSU , rCSU
             (CSU does not take into account chemical properties of atoms)
rCSU       - net contact from rCSU
Count      - number of contacts between residues
MODEL      - model number

      ID     I1  AA  C I(PDB)     I2  AA  C I(PDB)        DCA       CMs     rCSU    Count Model
==========================================================================================
R      1      1  MET A    1        2  GLN A    2      3.8045     1 1 1 0     -8      363     0
R      2      1  MET A    1        3  ILE A    3      6.5762     1 1 1 1     84       84     0
R      3      1  MET A    1       16  GLU A   16      5.5057     1 1 0 0    -13       32     0
R      4      1  MET A    1       17  VAL A   17      5.3638     1 1 1 1    246      297     0
R      5      1  MET A    1       18  GLU A   18      5.8910     0 1 1 1     70       78     0
R      6      1  MET A    1       19  PRO A   19      7.9600     1 1 1 1     66      132     0
R      7      1  MET A    1       56  LEU A   56     11.1189     1 1 1 1      1        1     0
R      8      1  MET A    1       61  ILE A   61     10.2682     1 1 1 0   -118      143     0
R      9      1  MET A    1       62  GLN A   62      8.1000     1 0 0 0      0        0     0
R     10      1  MET A    1       63  LYS A   63      5.3906     1 1 1 1     59      150     0
R     11      2  GLN A    2        1  MET A    1      3.8045     1 1 0 0     -1      350     0
R     12      2  GLN A    2        3  ILE A    3      3.7956     1 1 1 0    -50      414     0
R     13      2  GLN A    2        4  PHE A    4      6.8977     1 1 1 0    -73      107     0
R     14      2  GLN A    2       14  THR A   14      6.6212     1 1 1 0     -7      114     0
R     15      2  GLN A    2       15  LEU A   15      5.6908     1 1 0 0    -18       31     0
R     16      2  GLN A    2       16  GLU A   16      4.6970     0 1 1 0     -4       54     0
R     17      2  GLN A    2       63  LYS A   63      5.7883     1 1 0 0     -8        8     0
R     18      2  GLN A    2       64  GLU A   64      5.5447     1 1 1 1    115      237     0
R     19      3  ILE A    3        1  MET A    1      6.5762     1 1 1 1    130      130     0
R     20      3  ILE A    3        2  GLN A    2      3.7956     1 1 0 0    -39      393     0
R     21      3  ILE A    3        4  PHE A    4      3.8196     1 1 1 0     -3      354     0
R     22      3  ILE A    3       14  THR A   14      5.8173     1 1 0 0      0       18     0
R     23      3  ILE A    3       15  LEU A   15      5.5135     1 1 1 1    210      210     0
```

### (B) SUPPORTING METHODS

#### *(B1) PH domain of phospholipase C δ1*

The CG model of the phospholipase C δ1 (PLCδ1) pleckstrin homology (PH) domain was built using the open-beta version of the Martini 3 force field and the crystal structure with the pdb code 1MAI [9]. The missing termini were modelled with the iTASSER server[10,11]. The protein was simulated with the Gō-like model as well as both EN models. For the Gō-like model, a total of 234 contacts were included (216 OV and 18 rCSU contacts). The depth of the LJ potential was $\varepsilon$ = 12.0 kJ/mol. Both EN types were tested in this case. We use the same cutoff distance of 0.8 nm for both of them, but different force constants of $k_{t1}$ = 700 kJ/(mol·nm$^2$) for the EN type 1 and $k_{t6}$ = 500 kJ/(mol·nm$^2$) for the EN type 6, respectively. In order to simulate the PLCδ1 PH domain, a peripheral membrane protein specifically binding to PI(4,5)$P_2$, a single PI(4,5)$P_2$ lipid was embedded in a POPC bilayer consisting of in total 706 lipids with a patch size of 15×15 nm$^2$. The phosphate groups of the inositol triphosphate in the crystal structure were fitted to the PO4 beads of CG PI(4,5)$P_2$ model. The system was neutralized and solvated in 0.15 M NaCl solution. The simulation box with a size of 15×15×14 nm$^3$ contained ~17,500 CG water beads representing ~70,000 water molecules, 189 $Na^+$, and 189 $Cl^-$ ions. Protein-membrane distance analyses and potential mean force (PMF) calculations were already detailed in previous work [12].

#### *(B2) T4 lysozyme L99A mutant*

A detailed description of the CG model of the T4 lysozyme L99A mutant and benzene is provided in a previous work[13]. In summary, the crystal structure[14] with the pdb code 181L was used as reference. The CG model was generated using the open-beta version of the Martini 3 force field[15,16]. Both Gō-like and EN models were generated. For the Gō-like model, a total of 355 contacts were included, with 333 OV and 22 rCSU. The depth of the LJ potential was $\varepsilon$ = 15.0 kJ/mol in this case. The EN type 6 model was set up using a force constant of 500 kJ/(mol·nm$^2$) and a distance cutoff of 0.9 nm. The system was solvated using the *insane.py* script[17], with a water solution with 0.15 M concentration of NaCl, mimicking physiological conditions. The final simulation box had dimensions of 10x10x10 nm$^3$ and contained ~8,850 CG water beads, representing ~35,400 water molecules, 93 $Na^+$, and 101 $Cl^-$ ions. One molecule of benzene was randomly placed in the solvent, which is equivalent to a benzene concentration in water of around 1.6 mM. Minimization and equilibration protocols were the same as used in our previous publication involving protein–ligand binding with Martini 3[13]. RMSF calculations were performed using a homemade Fortran program, based on MDLovoFit code[18]. As a standard reference for the trajectory alignment, we used the average structure obtained in the CG simulation of each system. Details of the other analyses performed, including ligand density and free energy estimates were already described in our previous work[13]. A total sampling time of 0.9 ms (30 independent trajectories of 30 μs each) per system was used for the analysis.

*(B3) Wild-type and G93A mutant of Copper, Zinc Superoxide Dismutase 1*

A detailed description of the CG models of Cu,Zn Superoxide Dismutase 1 (SOD1) is given in a previous work[19]. In brief, an equilibrated atomistic structure of the SOD1 monomer, based on the crystal structure with the pdb code 2C9V, was used as the atomistic reference structure[20]. The CG model of SOD1 was built using the open-beta version of the Martini 3 force field[15,16]. In addition to the Gō-like model, monomeric SOD1 was simulated with both EN models, too. For the Gō-like model, a total of 331 contacts were included (301 OV and 30 rCSU contacts). The depth of the LJ potential was $\varepsilon$ = 12.0 kJ/mol. The EN models contained 542 harmonic bonds using a distance cutoff of 0.8 nm. The soluble protein SOD1 was neutralized and solvated in 0.15 M NaCl solution. The simulation box with a size of 8×8×8 $nm^3$ contained ~4,600 CG water beads representing ~18,400 water molecules, 54 $Na^+$, and 48 $Cl^-$ ions. The CG models and simulation boxes for wild-type and G93A mutants are the same, except for the mutation site, which in the Martini 3 open-beta model corresponds to a change from a SP1 to a P1 bead. Details of the minimization and equilibration procedure, and also analyses performed (including the RMSF calculations and the integrated absolute difference in the distance distributions) were already described previously[19]. A total of twelve 40 μs production runs were used for the analysis, with a total sampling of 480 μs per system.

*(B4) Nanomechanics of protein complexes by GōMartini model*

**XMod-Doc:Coh complex**

The system under study was the XMod-Doc:Coh complex, which is part of the bacterial adhesion protein complex in *Ruminococcus flavefaciens* (*R.f.*) and responsible for the recognition and attachment towards cellulose microfibrils. This system was built using the Martini 3 force field and the crystal structure with PDB ID: 4IU3[21]. The central part of this protein complex consists of the cohesin-dockerin complex (hereafter denoted by Coh:Doc) and dockerin protein is followed by the X-module (i.e. XMod) which is the anchored protein (see Fig. 5A). Importantly, this protein complex includes three $Ca^{2+}$ ions close to the protein-protein interface which are coordinated by negatively charged aspartates. In order to maintain a stable coordination sphere, the $Ca^{2+}$ ions were bonded to their respective aspartate neighbors, using a harmonic potential with a force constant of $k_b$ = 7,500 kJ/(mol·nm$^2$) and minimum distances according to the crystal structure. The CG Martini 3 representation is depicted in Fig. 5A. For the GōMartini model, a total of 1036 contacts were included using the OV and rCSU contact map determination. The interface of the coh-doc complex was represented by 59 contacts. The depth of the LJ potential in the GōMartini study was set equal to $\varepsilon$ = 9.414 kJ/mol (as in the ref. [22]). This protein complex was placed in a simulation box of 16×16×95 nm$^3$ containing 0.15 M NaCl solution. The number of CG water molecules was ~216,000 representing ~864,000 water molecules. The extended length in the z-dimension enabled the protein pulling along this direction. The pre-production system was equilibrated following a standard protocol. First, the system was energy minimized via steepest descent during 5,000 steps, then an NVT ensemble at 300 K was used to equilibrate the temperature, and finally, an isotropic NPT ensemble at 300 K and 1 bar to achieve ambient conditions.

Nanomechanical studies were carried out via steered molecular dynamics (SMD) simulations at constant speed, see simulation box in Fig. 5(B). The SMD protocol in GROMACS is based on a modified umbrella center-of-mass pulling protocol. The vector defined between the N and C termini of the XMod-Doc:Coh complex was aligned with respect to the longest axis of the simulation box. To reproduce experiments conducted by single-molecule force spectroscopy with the atomic force microscope (AFM-SMFS) [23], the coordinates of the cohesin C-terminal residue (i.e. GLY-201) were immobilized, as such condition satisfies the anchoring of the cohesin on the cantilever by a covalent bond. On the other side of the complex, the N-terminus (ASN-5) located at the X-Doc domain is coupled by virtual particles and pulled via a harmonic potential along the main box axis. It is important to note that all CG beads in each pulling residue are considered necessary to perform the pulling simulation. The pulling rate and spring constant correspond to $5x10^{-4}$ nm/ps and 37.6 kJ/(mol·nm$^2$) respectively. The latter parameter corresponds to a typical experimental AFM cantilever stiffness[6]. The ensemble of pulling simulations comprises a set of 100 replicas.

**RBD:H11-H4 complex**

Here we study a protein complex system that involves the interaction between a single-domain antibody (i.e. nanobody) and the receptor-binding domain (RBD) portion of the SARS-CoV-2 spike protein. In this regard, the nanobody named H11-H4 and the RBD form a mechanostable protein complex with PDB ID: 6ZH9. The entire system was modeled by the Martini 3 force field. The same protocol as for the XMod-Doc:Coh complex was employed. The GōMartini model requires a total of 715 contacts: 404 contacts within the RDB, 285 within the H11-H4, and 26 between the two proteins. The dimensions of the water box for the RBD:H11-H4 complex were 16x12x90 $nm^3$ containing 0.15 M NaCl solution. There were 135,669 CG water beads representing 542,676 water molecules. To conduct the nanomechanical studies at constant pulling speed and avoid instability in SMD protocol, we applied constraints along the z-axis on the RBD residues GLY-526, PRO-527, and LYS-528 and similarly in x- and y-axis on the residues SER-126, SER-127, and LYS-128 in H11-H4. The value of the dissociation energy of the LJ potentials was set to $\varepsilon_{LJ}$ = 15.0 kJ/mol. The position of RBD residue LYS-528 was immobilized and the coordinates of LYS-128 in H11-H4 were chosen for SMD simulation. Nanomechanical studies for this complex by all-atom MD simulation are available in the literature, and they report very different values for the rupture force associated with this protein complex. For instance, restraints were applied to the backbone atoms of the RBD with a spring contact of $k_b$ =1000 kJ/(mol·$nm^2$)[24,25] to prevent large deformations in the RBD. Also, the residue in the SMD protocol was pulled with velocity equal to 5x$10^{-4}$ nm/ps and coupled to another harmonic potential with a spring constant equal to 600 kJ/(mol·nm2). The other study fixed the position of certain residues located at the RBD interface with H11-H4 and steered the position of other residues (located in H11-H4 interface with RBD) by SMD protocol with a spring constant of 60 kJ/(mol·nm2) and the speed of $10^{-5}$ nm/ps[24,25]. We employed unrestrained GōMartini simulations[6] to validate both approaches and thus to assess the impact of restraints on the nanomechanics of protein complexes. We performed 100 replicas for each speed and SMD simulation.

*(B5) Protein flexibility benchmarks*

We estimated protein flexibility for a set of 4 soluble proteins (ubiquitin, titin, cohesin, and Man5B with PDB ID: 1UBQ[26], 1TIT[27], 1AOH[28], and 3W0K[29], respectively) and 2 transmembrane proteins (human AQP1 with PDB ID: 4CSK[30] and yeast Ist2). For Ist2, no high-resolution structure is available in the PDB, therefore homology modeling techniques were employed to build a 3D model, limited to the transmembrane domain (TMD) (residues 70-589, retrieved through the UniProtKB database, http://www.uniprot.org/, accession IDs: P38250). We modeled Ist2 TMD using the trRosetta web server with a fully automated structure prediction pipeline. The similarity of the model to the experimental closed conformation nhTMEM16 (PDB ID: 6QMB) was quantified using the global superposition metric template modeling score (TM-score), calculated using TM-align[31], which yields scores from 0 (arbitrarily different) to 1 (identical structures). In the case of the Ist2-TMD, the TM-score of the model was 0.78. MODELER v10.1[32] was then used to build the homodimer model of the closed state of the protein, using the PDB ID: 6QMB as template. One hundred models were generated, and the one with the lowest discrete optimized protein energy (DOPE) score was adopted for the MD simulations.

To set up all-atom simulations, soluble proteins were solvated in a cubic simulation box of CHARMM TIP3P water molecules[33,34], and the final dimensions of the box were ~ 7 × 7 × 7 nm. The structures of the AQP1 and Ist2 were embedded respectively into pure POPC and a lipid mixture of DYPC, DYPE, and DYPS bilayers. The interactions were described by the CHARMM36m force field[35]. TIP3P water model[33,34] and 0.15 M KCl were added to solvate and neutralize the membrane-protein system. The AA simulations systems were setup using the CHARMM-GUI[36,37].

CG simulations of analogous systems (i.e., same protein, same lipids), were set up using the *insane.py* script. As mentioned in the previous sections, to generate a GōMartini model for protein it is necessary to use *Martinize2* and *create_goVirt.py* script. The contact map of all atom structure was obtained from the http://info.ifpan.edu.pl/~rcsu/rcsu/index.html web server using default settings for the radii and Fibonacci number, and was based on a single high-resolution structure. After establishing the initial network of bonds, we explored the effect of varying dissociation energy (ε) for the Lennard-Jones (LJ) potentials by scanning a range of ε values from 12 to 27 kJ/mol for each protein, with three independent simulations of 250 ns each, for each protein. The root mean square fluctuation (RMSF) of particle positions was then used as a simple measure of protein flexibility, and compared between the different CG simulations (with different values of ε) and the reference AA simulation. It is worth mentioning that we assumed a conversion factor of 4 for the effective time of the simulations to compare the CG and AA trajectories. Therefore, we compared the RMSF obtained in 1 μs of atomistic simulation with 250 ns of coarse-grained simulation. The CG setup providing an RMSF most similar to the AA reference was selected as the optimal one, reproducing the overall flexibility of these proteins observed in AA simulations of comparable duration. The optimal ε values were different for the 6 proteins analyzed: ε = 18 kJ/mol for 3W0K, ε = 15 kJ/mol for 1UBQ and 1AOH, ε = 25 kJ/mol for 1TIT, ε = 11 kJ/mol for Ist2, and ε = 12 kJ/mol for AQP1.

While this procedure provided overall reasonable agreement with AA simulations in terms of protein dynamics, the flexibility of some specific regions of the CG models remained underestimated. To further improve the performance of the model, we found that it was necessary to selectively omit certain bonds in the Go network (i.e., defined between the virtual Gō beads), particularly in regions of higher flexibility. We then devised an automatic procedure to select the bonds to be discarded, based on the contact frequencies of each residue in the AA simulation. To this end, we used weighted residue contact maps as calculated by the MD-TASK script[38], that determines contact frequencies for each residue with neighboring residues during an MD simulation. In a residue interaction network, each residue in the protein is a node in the network; an edge between two nodes exists if the Cβ atoms (Cα for Glycine) of the residues are within a user-defined cut-off distance from each other. To find a cut-off which reproduces protein dynamics as observed in AA simulations, we calculated the contact frequencies of residues by scanning the cut-off in the range 0.75 – 1.0 nm for each protein, and extracted the residue contacts in the AA trajectories. We employed an in-house python script to extract residue contact data and edit the existing itp files. If the contact frequency for a specific node in AA simulation is low, the corresponding Gō bond is removed in the itp file. These modifications were applied to both *_exclusions_VirtGoSites.itp and *_go-table_VirtGoSites.itp files (initially generated by the create_goVirt.py script).

**Simulation parameters**: energy minimizations were carried out with the steepest descent algorithm for 50,000 steps. Subsequently, two equilibration phases, each spanning 250 ns, were performed, using the leap-frog algorithm. Throughout these equilibration phases, position restraints (k = 1000 $mol^{-1}$ $nm^{-2}$ and 100 kJ $mol^{-1}$ $nm^{-2}$, respectively) were applied exclusively to the backbone atoms of the protein structure. After the equilibration steps, unbiased MD simulations were performed. All systems were simulated under periodic boundaries in the NPT ensemble. The temperature and pressure were controlled with a velocity-rescale thermostat and a Parrinello–Rahman barostat, respectively[39]. Solvent and solute (lipids and protein) were independently coupled to a temperature and a pressure bath during the all simulations. Each system was replicated and assigned with different initial velocities to generate independent simulations. Snapshots were sampled every 25 ps in CG and every 100 ps in AA simulations. Details of parameters and setup of all simulated systems are provided in Table S2.

**Table S2.** Details of simulation systems.

| PDB ID | Residues | #Atoms in AA model | AA simulation time (μs) * | #Particles in CG model | total contacts OV+rCSU | ε | Weight Edge | Cut off (Å) | Percentage of ignored Gō bonds |
|---|---|---|---|---|---|---|---|---|---|

| | | | | | | | | | |
|---|---|---|---|---|---|---|---|---|---|
| 1UBQ | 76 | 24597 | 2×1 | 2212 | 304 | 15 | 0.60 | 9.0 | 3.94 % |
| 1TIT | 89 | 29313 | 2×1 | 2759 | 384 | 25 | 0.95 | 9.5 | 9.8 % |
| 1AOH | 147 | 34894 | 2×1 | 3453 | 705 | 15 | 0.95 | 9.0 | 12.82 % |
| 3W0K | 330 | 44327 | 2×1 | 4372 | 1644 | 18 | 0.95 | 10.0 | 13.7 % |
| 4CSK | 932 | 111567 | 3×1 | 10470 | 954** | 12 | 0.65 | 7.5 | 14.37 % |
| Ist2 | 1040 | 697564 | 2×1 | 58735 | 2122** | 11 | 0.75 | 10.0 | 25.33 % |

* CG data saved every 25 ps – AA data saved every 100 ps

** per chain

*(B6) Intrinsically disordered proteins*

Initial conformational ensembles of 10000 IDPs from the set of Thomasen et. al.[40] were generated using flexible meccano through the nmrbox server[41], and side chain reconstruction was performed using pulchra, as described in the protocol of Ahmed et al[42]. From these ensembles, a conformer was chosen from the 95th percentile of the radius of gyration distribution.

For atomistic reference simulations, each protein was centered in a dodecahedral box 3 nm away from the sides, solvated, and had the salt concentration fixed to the experimental reference value. Simulations were then conducted using the CHARMM36m force field[35], with the CHARMM36m TIP3P water model[33,34], as recommended for simulations of disordered proteins. Standard simulation settings for the CHARMM36m force field were used: long range electrostatics were calculated using PME with a cutoff of 1.2 nm, LJ interactions were treated with a cutoff of 1.2 nm and a force switch modifier from 1 nm. After the addition of salt at the experimentally relevant concentration, the system was energy minimized. The systems were then equilibrated in the NVT and NPT ensembles for 1 ns each using a timestep of 2 fs. The simulation temperature was set to the temperature of the respective experimental temperature of the measurement of the IDP's radius of gyration. Temperature was maintained using the velocity-rescaling thermostat, and pressure at 1 bar was maintained using the Berendsen[43] and Parrinello-Rahman[39] barostats for equilibration and production respectively. Subsequently, production simulations were carried out for 100 ns also using a timestep of 2 fs. The trajectories were subsequently mapped to Martini resolution using fast_forward, a recently developed python library for this purpose. The mapped trajectories were then used for analyzing bonded distributions of backbone dihedrals and sidechain dihedrals as detailed in the main text.

For CG simulations using the Martini 3 force field, we firstly generated CG parameters and coordinates using the *Martinize2* program.[44] The initial parameters were then adapted using custom scripts based on the Vermouth library, introducing backbone dihedral potentials to reproduce the ones observed from atomistic simulations, and adding virtual Gō sites as appropriate. Side chain parameters were fixed using the -scfix flag in *Martinize2*[44]. Simulations were set up similarly to the atomistic ones: the conformer was placed in a dodecahedral box 3 nm from the edges, solvated and had the salt concentration and temperature fixed at the corresponding experimental ones. Production simulations were run for 10 μs using the standard Martini simulation parameters described above.

*(B7) Biomolecular condensates*

**Artificial IDPs**

Parameters of the aIDPs designed by Dzuricky *et al.* were generated using the optimized Martini IDP model described. Initial configurations of 50 aIDPs were generated in a 30x30x30 nm cubic box using *Polyply* and solvated to result in a system at 5% protein weight. Salt was added to a concentration of 0.2 M. Production runs were simulated for 5 μs.

**Short peptide synthon**

The short peptide synthon consists of a disulfide moiety linking two FL dipeptides via the C termini (FLssLF). The two terminal dipeptides were coarse grained with coil secondary structure, using the Martini 3.0 force field. The linker related bonded parameters were optimized against atomistic reference simulation. We included virtual Gō sites to explore different additional interaction strengths between the Gō and water beads (106%: $\sigma = 0.465$ nm, $\varepsilon = 0.2598$ kJ/mol; 108%: $\sigma = 0.465$ nm, $\varepsilon = 0.3464$ kJ/mol ).

Due to lack of an experimental pKa value, the pKa of the terminal amine was predicted by the MOLGPKA and Graph-pKa web servers, with a mean value of 6.84[45,46]. Mimicking experimental conditions of pH=8, 36 ionized and 528 neutral species were placed in a 8x8x90 nm rectangular box (~110 mg/ml) to keep the simulation tractable and solvated using regular Martini 3 water beads. To reduce the simulation length required for convergence, a slab-like initial configuration was used for simulation following the same protocol in [47], in which pure peptides were squeezed to slab-like and then solvated into the rectangular box. The equilibration was conducted for 100 ns in the NPT ensemble at 298K and 1 bar, maintained by the velocity-rescale thermostat and the Berendsen barostat[43] (with coupling constant 4 ps), respectively. A production run was conducted for 3 µs under the same conditions, except switching to the Parrinello-Rahman barostat (with coupling constant 12 ps). During both phases, semi-isotropic pressure coupling was imposed to allow the system to equilibrate only along the z-axis. The Verlet cut-off scheme was used to treat non-bonded interactions with a Van der Waals interaction cut-off of 1.1 nm. The reaction-field method was used to treat Coulomb interactions using a 1.1 nm cut-off with a dielectric constant of 15. The mass density profile was computed by gmx density, omitting the first 1 µs of the final trajectory.

*(B8) Transmembrane peptides*

Transmembrane WALP simulations were prepared as described elsewhere[48]. In short, Martini 3 topologies were created from ideal $\alpha$-helical transmembrane WALP structures, created from their sequence using Avogadro[49]. As WALPs are capped on both termini, with N-terminal acetylation and C-terminal n-methyl amidation[50], and neither capping group has been explicitly parameterized for Martini 3, the terminals were modeled by assigning a non-charged backbone particle type (P2). The peptides were aligned vertically within a 10×10 $nm^2$ dimirystoylphosphatidylcholine (DMPC) membrane in ideal transmembrane configurations, with a 9 nm simulation box height, built using the *insane.py* script[17]. The system was then solvated with Martini 3 water beads, and a NaCl ionic strength of 0.15 M was added. For each simulation, 5 replicas of at least 30 µs were run. To facilitate the observation of WALP ejection from the membrane, a temperature of 310 K was used.

*(B9) RAD16-I peptide*

The RAD16-I peptide structure was built using the PyMol software[51]. The assembly of the 2-strand antiparallel beta-sheet as well as correct side-chain orientation was obtained through an in-house developed Python script, using MDAnalysis and numpy packages. Three CG systems were then created: a beta-sheet with regular *Martinize2* structure; beta-sheet with inter-chain Gō model; and a beta-sheet structure with interaction bias between virtual Gō sites and water beads. To generate each CG model, we made use of *Martinize2*[44], whilst the create_goVirt.py script was used additionally to apply the Gō model and water interaction bias. To ensure the peptide strands did not curl into themselves, the dihedrals and BBB angles were restrained at 180º and 137º, respectively.

All systems were solvated, in a 9x9x6 dodecahedron box, with regular Martini 3 water beads, and neutralized with Na+ cations and Cl- anions. For every simulation, the nonbonded interactions had a cut-off distance of 1.1 nm and we used reaction-field electrostatics with a dielectric constant of 15 and an infinite reaction-field dielectric constant to treat the Coulombic interactions. The Verlet list scheme was used to update the particle neighbor list. We made use of a V-rescale thermostat with a coupling time of 3.0 ps to maintain the temperature at 310 K. An isotropic constant pressure was used for all simulations. They were coupled to 1.0 bar using the Parrinello-Rahman barostat with a relaxation time of 12.0 ps. The simulations were run at a 20 fs time-step, succeeding the initial energy minimization along with the temperature and pressure equilibration runs. For each system, 3 replicas were run for at least 10 μs.

### (C) SUPPORTING RESULTS

#### *(C1) PLCδ1 PH domain: PMFs for strong lipid binding*

Figure S3 shows the protein-membrane distance of the center of mass of the PLCδ1 PH domain and the PI(4,5)$P_2$ headgroup. For each protein model, 10 replicas of 2 μs each were performed.

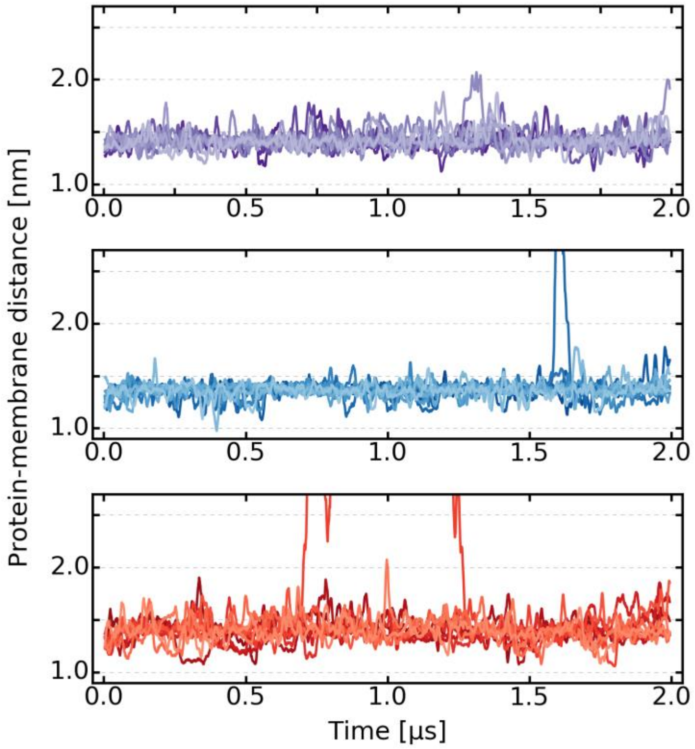


**Figure S2. Protein-membrane distance of PLCδ1 PH domain bound to a PI(4,5)$P_2$ lipid.** Distance between the PI(4,5)$P_2$ headgroup and the center of mass of the PLCδ1 PH domain. The protein was modeled with an EN model type 6, 500 kJ/(mol $nm^2$), and a cutoff of 0.8 nm (top), an EN model type 1, 700 kJ/(mol $nm^2$), and a cutoff of 0.8 nm (middle), and a Gō-like model with an $\varepsilon_{LJ}$ = 12.0 kJ/mol (bottom). For each protein model, 10 replicas of 2 μs each are depicted.

Figure S4 shows the effect of excluding non-bonded interactions for the EN bonds using a weak force constant of 500 kJ/(mol $nm^2$). Excluding the non-bonded interactions drastically increases the depth of the PMF by about 13 kJ/mol from a minimum at −21.1 kJ/mol (bond type 6) to −34.0 kJ/mol (bond type 1). This reinforces the recent observation that too low force constants can induce too short bond lengths which contributed to the overestimation of protein-protein interactions in Martini 2 with elastic network models[52].

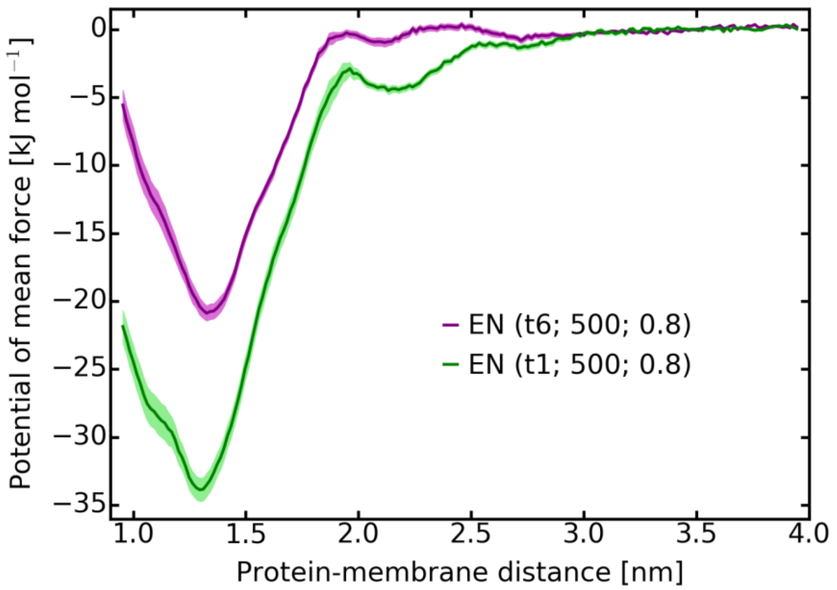


**Figure S3. Potentials of mean force of the membrane binding of PLCδ1 PH domain using different bond types of the elastic network model.** The protein was modeled with an EN in both cases using a force constant of 500 kJ/(mol nm$^2$) and a cutoff of 0.8 nm. The only difference is the exclusion of non-bonded interactions (bond type 1 in GROMACS, green) and not excluding non-bonded interactions (bond type 6, violet).

*(C2) Assessing the mechanic stability of XMod-Coh:Doc complex from bacteria*

The improved GōMartini implementation can also be applied to study the nanomechanical stability of protein complexes, in this case, via SMD simulations at constant speed mode. These *in silico* experiments can help interpret AFM-SMFS experiments at moderate cantilever speeds. The case study is a fundamental part of the extracellular multienzyme system known as the cellulosome[53]. The full-length cellulosome is secreted by bacteria and generally each protein component self-assembles extracellularly in anaerobic thermophilic conditions. The cellulosome is considered a very advanced nanomachine, responsible for efficiently dismantling the plant cell walls into their monomeric subunits. We consider the XMod-Doc:CohE complex. This system is formed by an Ig-like X-Module (XMod) adjacent to the scaffold-borne Dockerin from the *R.f.* cellulosome CttA protein, and the cell-wall anchored Cohesin-E (CohE) (see Figure S5A). The native geometry of this protein complex was characterized by AMF-SMFS studies and discovered its resilience to withstand pulling forces in the range of 500-800 pN at loading rates ranging from 2-300 nN/s[54]. Note that AFM-SMFS experiments are performed at several orders of magnitude much smaller speed (e.g. 6.4 μm $s^{-1}$) in comparison to SMD simulations. We performed GōMartini simulations by SMD (see Figure S5B), and identified two main pathways that led to the dissociation of the protein complex. We show the intermediary steps behind the one-step dissociation process (see Fig. 5C), which in the F-displacement plot is represented by one single force peak. The process of dissociation starts by the stretching of XMod-Doc at distances of 14 nm, at which point we observe the maximum applied forces of 600-700 pN. At a large distance of ~16.5 nm, we observe that XMod-Doc is stable enough to disassociate from Dockerin, which is further destabilized and starts losing Gō contacts at the Doc:Coh interface. Complete rupture occurs at distances larger than 17 nm. Another type of process identified by our simulation corresponds to a three-step dissociation process (see Figure S5C). Here the process of dissociation starts by the unfolding of XMod at distances from ~15 nm up to 36 nm. As in the one-step process, XMod-Doc is pulled away slightly from the Dockerin, but the XMod should first undergo complete unfolding before the interface of Doc:Coh can be dissociated. The reported maximum forces are similar to the one-step process (i.e. 600-700 pN). The following two peaks in the Force-displacement curves are associated with the unfolding process of XMod-Doc. After the complete unfolding of XMod, the force registered by our simulation was 300 pN and the Coh:Doc interface remained stable. A similar mechanism of dissociation for this complex was reported by AFM-

SMFS, where a bimodal distribution of maximum applied forces was observed, with high peaks close to 550 pN and 700 pN.

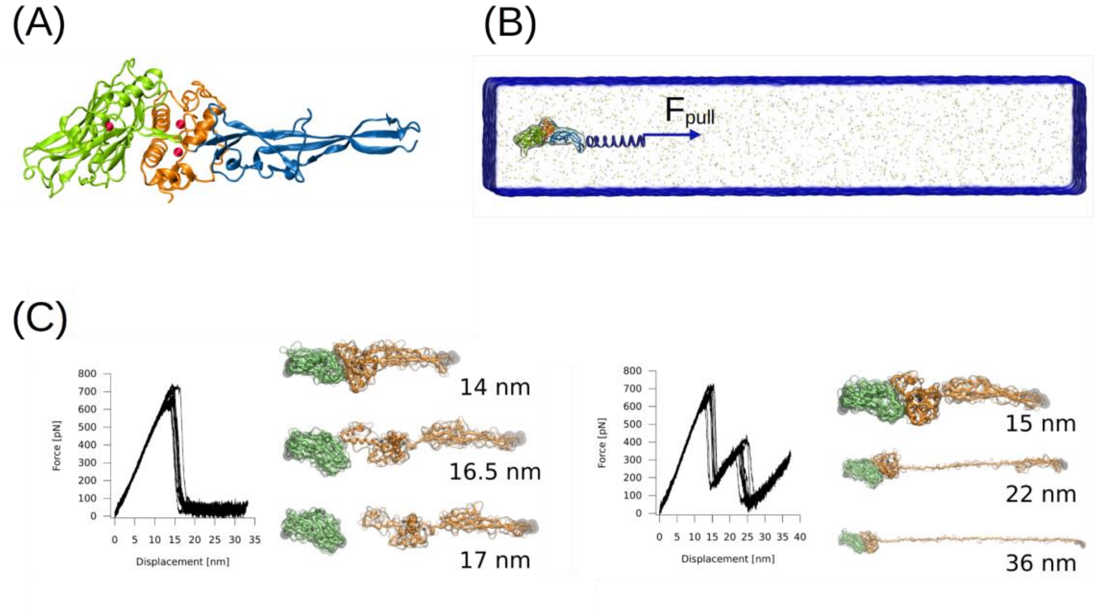


**Figure S4. Nanomechanics of Doc:Coh complex by GōMartini simulations.** (A) Structure of the XMod-Doc:Coh complex (PDB: 4IU3). Cohesin, Dockerin and X-Module are represented in green, orange, and blue, respectively. $Ca^{2+}$ atoms are represented by red spheres. (B) Simulation box of the protein complex in solution. The pulling direction along the long axis of the protein complex is also represented. (C) Force-displacement profiles obtained from the GōMartini SMD simulations of the *R.f.* cellulosome CttA protein. Force profiles show two important dissociation processes reported by all-atom MD simulation and AFM-SMFS experiments[23]. Left panel depicts the one-step or single peak event that is induced by the detachment of the XMod-Doc from Coh, whereas the right panel shows the corresponding profiles associated with the unfolding of XMod.

*(C3) Improving contact maps and strength of interactions*

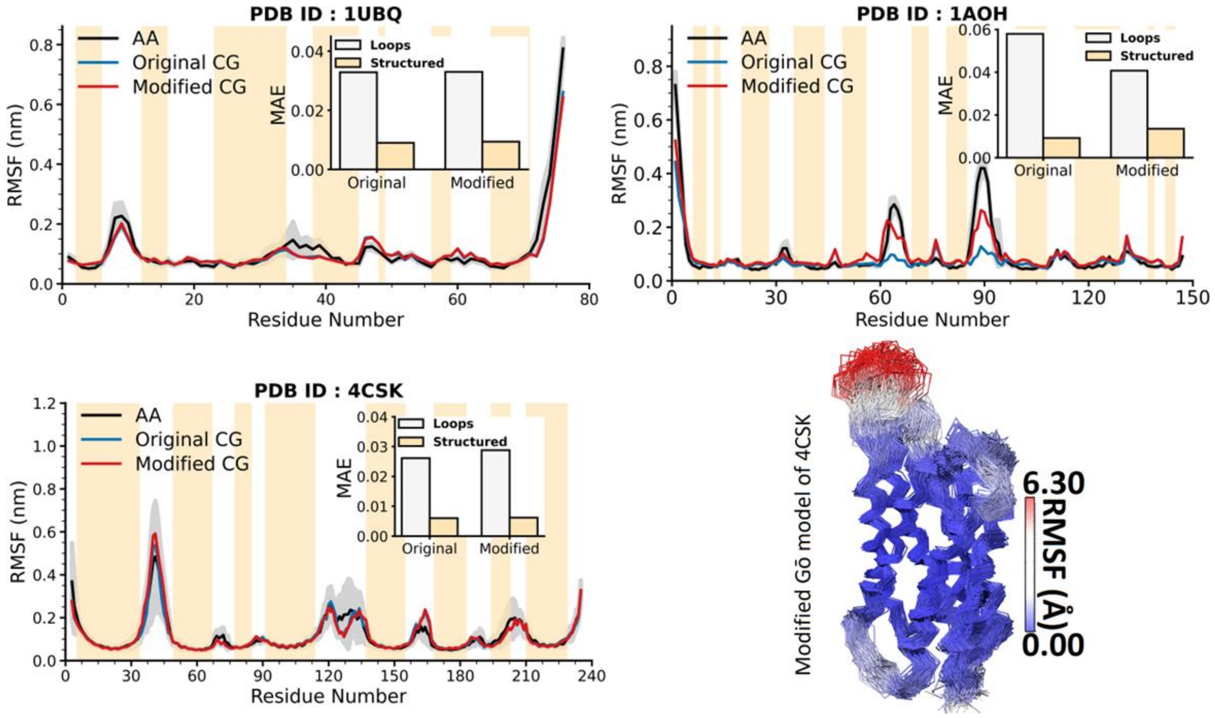


**Figure S5**. Residue RMSF Comparison between GōMartini and Atomistic Simulations for three proteins (1UBQ, 1AOH, and 4CSK) with Mean Absolute Error (MAE) Illustration.The bottom-right panel presents the flexibility of the protein backbone beads (BB) during simulations using the modified GōMartini model for one chain of the protein 4CSK (AQP).

*(C4) IDPs and biomolecular coacervates*

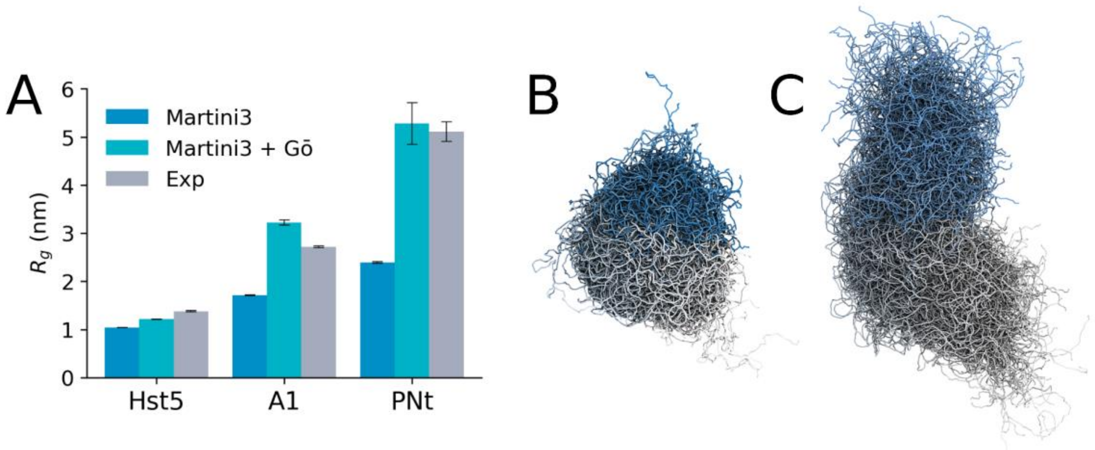


**Figure S6. Preliminary results testing of increased protein-water interactions in Martini IDPs**. A) Comparison of the $R_g$ of a subset of three IDPs from the reference set of Thomasen et al[40], comparing native Martini3, Martini3 + Go potentials of ε = 0.5 kJ/mol, and experimental data. B), C) Illustration of the expanded ensemble of PNt without (B) and with (C) increased protein-water interactions.

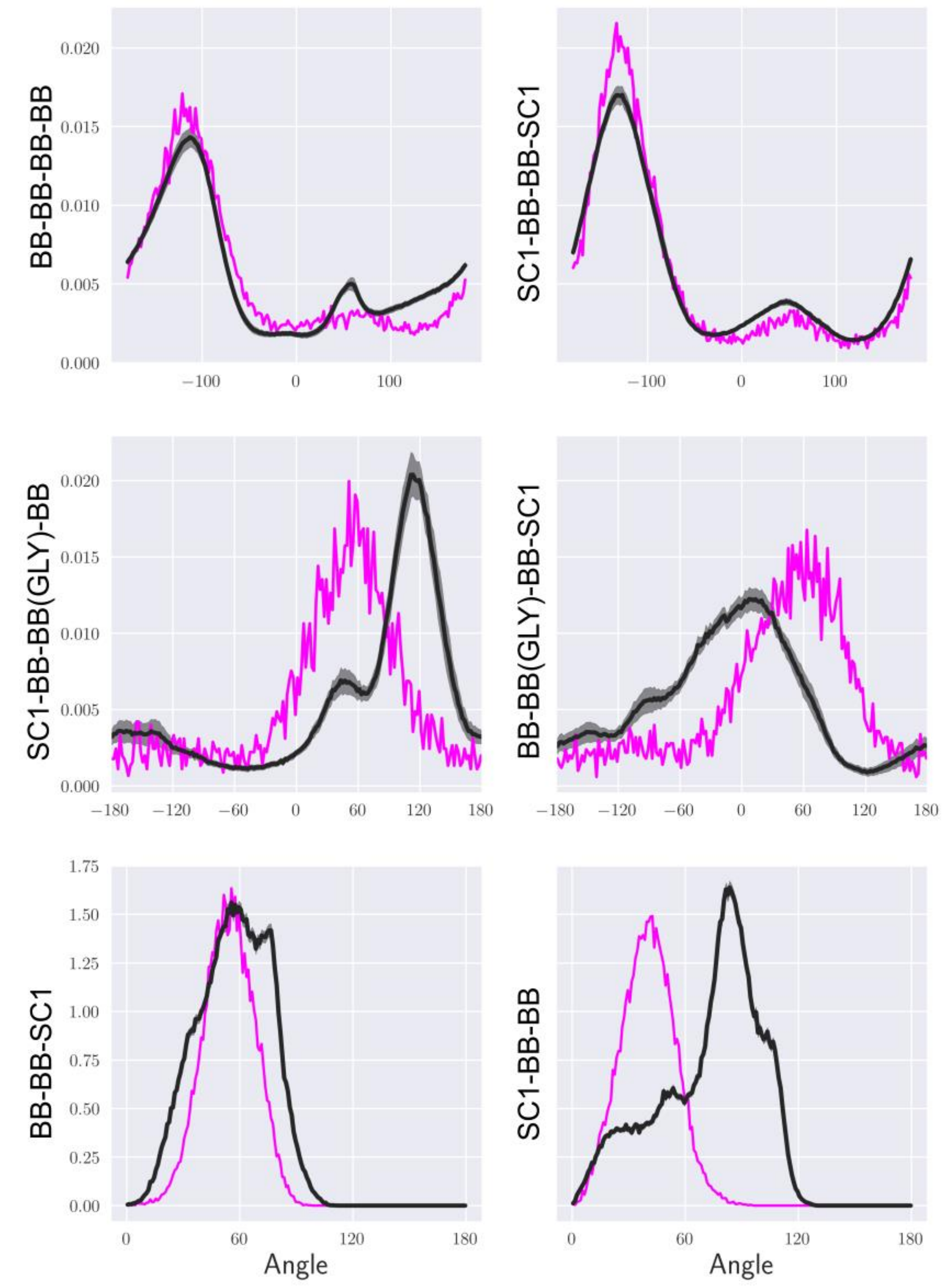


**Figure S7. Application of additional bonded potentials to IDPs**. Distributions from mapped atomistic simulations (black) of the reference set of Thomasen et al[40] were matched by the addition of extra bonded potentials in Martini models (pink).

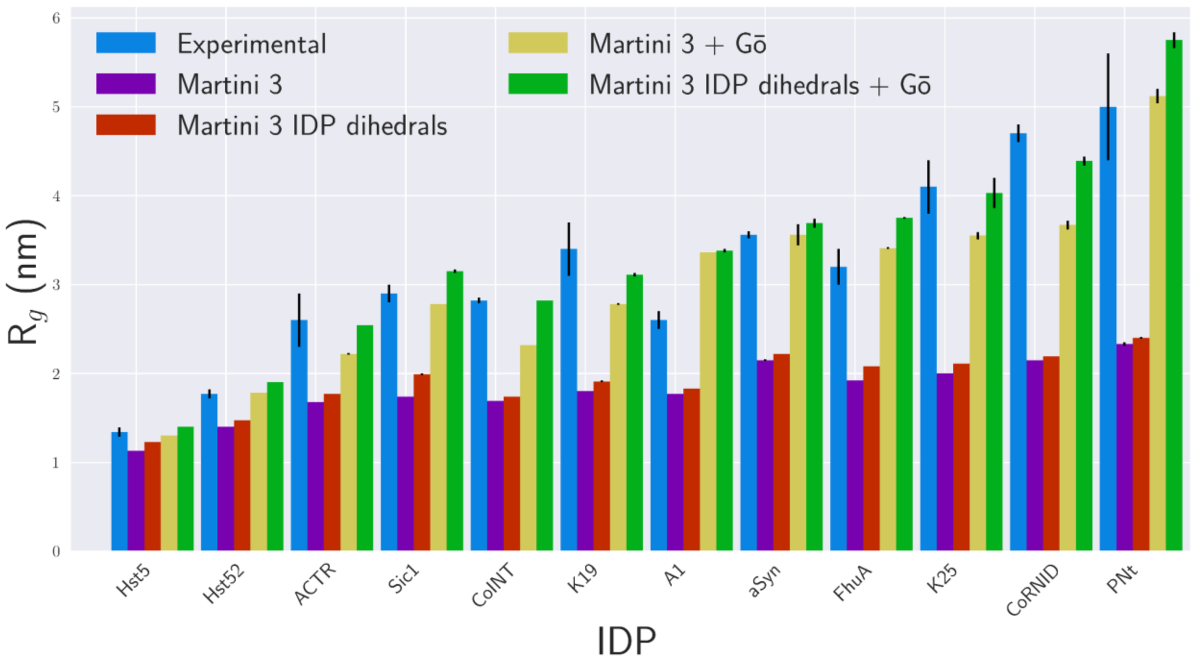


**Figure S8**. **Radii of gyration of different models of IDPs**. The target set of IDPs of Thomasen et al.[40] simulated with iterations of the Martini IDP model. The experimental reference is shown in blue. The default Martini 3 parameters are shown in Purple. The model with the default Martini 3 non-bonded parameters, and the IDP dihedral potentials from figure S7 are shown in Red. The default model without further bonded parameters, but with the addition of the virtual Gō site, are in yellow. The final model with both additional bonded and non-bonded parameters is in green. The blue, purple, and green bars are shown in Figure 8 of the main text. The mean absolute error across each set is 1.35, 1.25, 0.36, and 0.28 nm respectively.

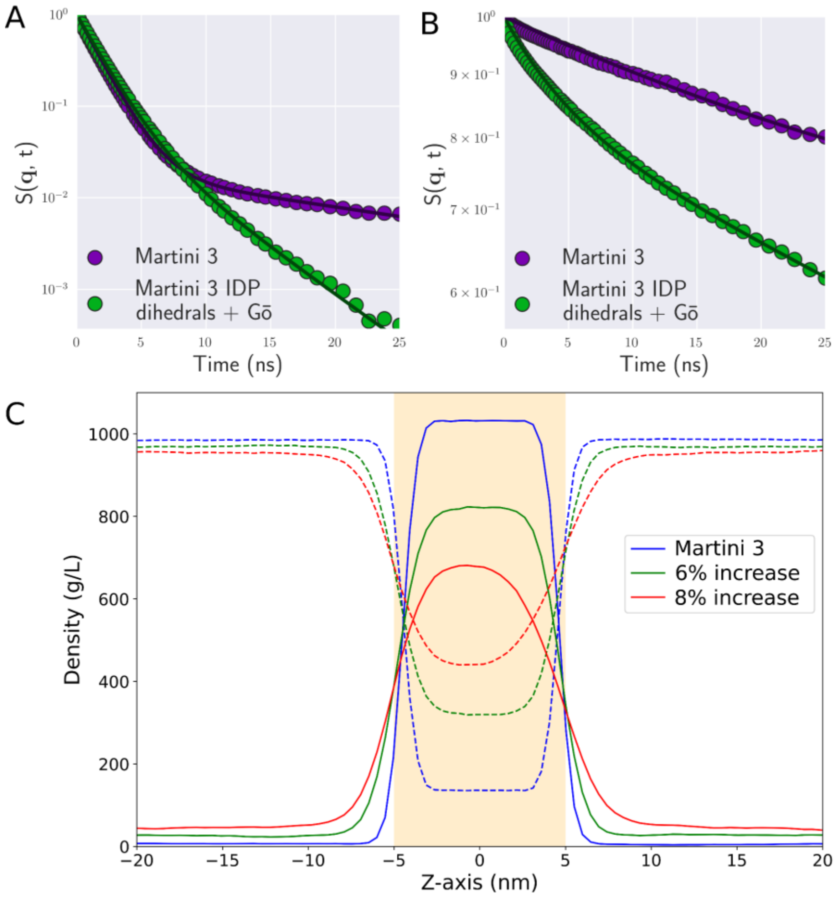


**Figure S9. Properties of biomolecular condensates.** A,B incoherent scattering functions of water and protein components of the two systems respectively, showing the different components measured for the default Martini aggregate (purple) and optimised IDP condensate (green). Incoherent scattering functions can be used to describe the diffusion coefficients in multiphase systems, through the fitting of weighted exponentials.[55] For the optimised system in green, there are two fitted components, indicating that both the proteins and water diffuse through two distinct environments, demonstrating the system is liquid-liquid phase separated. Further, for the water component of the system, there are still clearly two components, but the slow component of the function has reduced significantly, from 0.75x10-12 $m^2$ $s^{-1}$ to 0.15x10-12 $m^2$ $s^{-1}$. This reduction suggests that within the protein-dense region of the system, the dynamics of the water have been significantly arrested, as a result of the aggregation of the protein. Indeed, the incoherent scattering curve measured on the protein component of the native Martini 3 system shows that while two exponential decay components were present in the optimized system, now only a single exponential decay for an aggregated dense phase can be observed. C) mass density profiles for the three peptide systems shown in Figure 8 of the main text, peptide density is shown with solid lines and water density with dashed lines.

*(C5) TM helices and beta-sheet peptides*

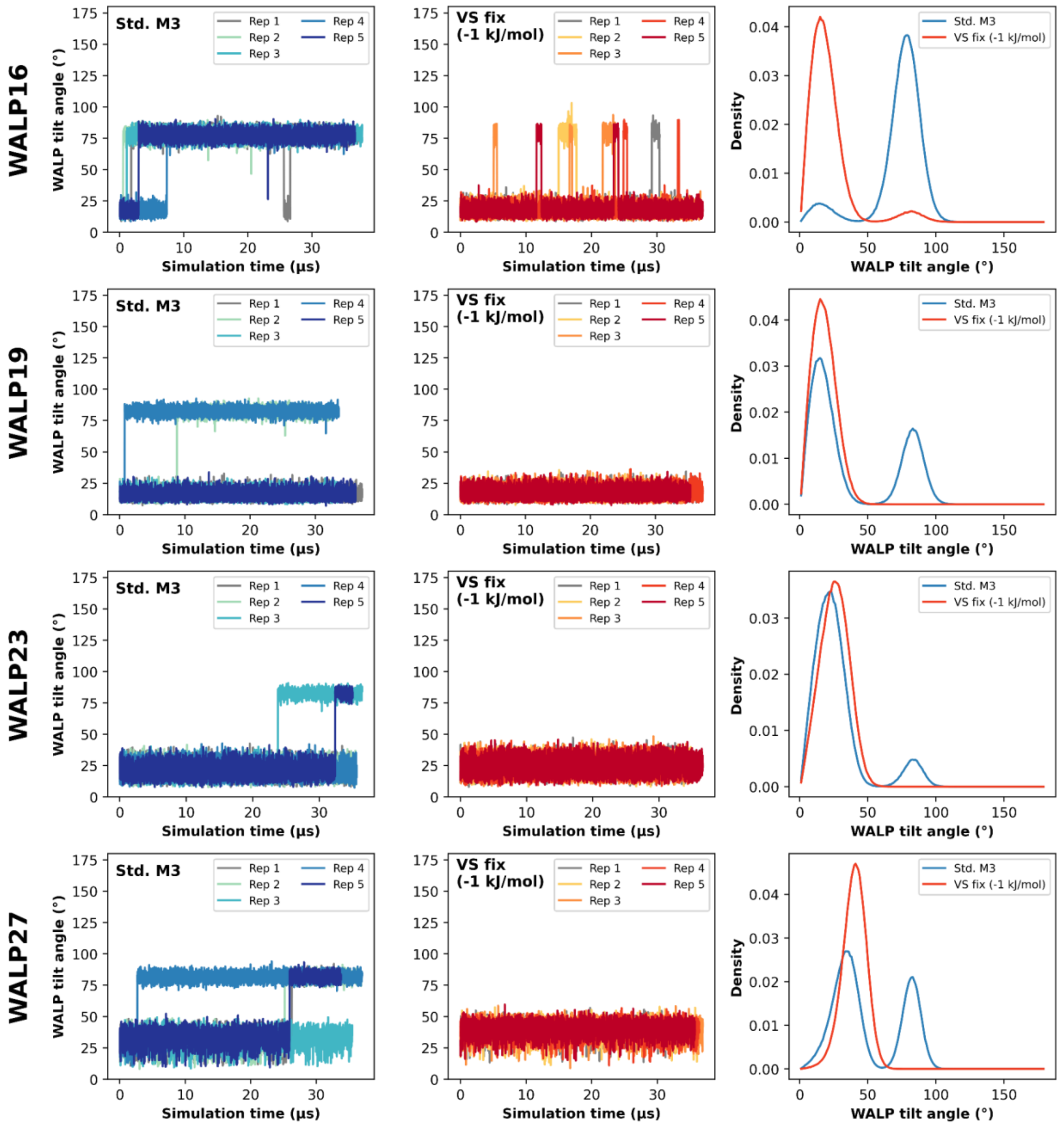


**Figure S10. Improving transmembrane WALP insertion.** Membrane inserted WALP peptide tilt angle distributions, for WALPs 16, 19, 23 and 27, recovered from simulations ran with our GoMartini implementation without any additional LJ interaction and with an additional LJ interaction between the virtual Gō sites and water beads with $\varepsilon$ = -1.0 kJ/mol. Both time traces of WALP tilt angle, and tilt angle distributions are shown.

(D) REFERENCES


1. Abraham, M. H., Chadha, H. S., Whiting, G. S. & Mitchell, R. C. Hydrogen bonding. 32. An analysis of water-octanol and water-alkane partitioning and the delta log P parameter of seiler. *J. Pharm. Sci.* **83**, 1085–1100 (1994).
2. Abraham, M. H., Platts, J. A., Hersey, A., Leo, A. J. & Taft, R. W. Correlation and estimation of gas-chloroform and water-chloroform partition coefficients by a linear free energy relationship method. *J. Pharm. Sci.* **88**, 670–679 (1999).
3. Natesan, S. *et al.* Structural determinants of drug partitioning in n-hexadecane/water system. *J. Chem. Inf. Model.* **53**, 1424–1435 (2013).
4. Sangster, J. Octanol-water partition coefficients of simple organic compounds. *J. Phys. Chem. Ref. Data* **18**, 1111–1229 (1989).
5. Hansch, C. & Leo, A. *Exploring QSAR.: Fundamentals and Applications in Chemistry and Biology*. (1995).
6. Liu, Z. *et al.* Mapping Mechanostable Pulling Geometries of a Therapeutic Anticalin/CTLA-4 Protein Complex. *Nano Lett.* **22**, 179–187 (2022).
7. Mahmood, M. I., Poma, A. B. & Okazaki, K.-I. Optimizing Gō-MARTINI Coarse-Grained Model for F-BAR Protein on Lipid Membrane. *Front Mol Biosci* **8**, 619381 (2021).
8. Wołek, K., Gómez-Sicilia, À. & Cieplak, M. Determination of contact maps in proteins: A combination of structural and chemical approaches. *J. Chem. Phys.* **143**, 243105 (2015).
9. Ferguson, K. M., Lemmon, M. A., Schlessinger, J. & Sigler, P. B. Structure of the high affinity complex of inositol trisphosphate with a phospholipase C pleckstrin homology domain. *Cell* **83**, 1037–1046 (1995).
10. Yang, J. & Zhang, Y. I-TASSER server: new development for protein structure and function predictions. *Nucleic Acids Res.* **43**, W174–81 (2015).
11. Yang, J. *et al.* The I-TASSER Suite: protein structure and function prediction. *Nat. Methods* **12**, 7–8 (2015).
12. Thallmair, V. *et al.* Two cooperative binding sites sensitize PI(4,5)P recognition by the tubby

domain. *Sci Adv* **8**, eabp9471 (2022).

13. Souza, P. C. T. *et al.* Protein-ligand binding with the coarse-grained Martini model. *Nat. Commun.* **11**, 3714 (2020).
14. Morton, A. & Matthews, B. W. SPECIFICITY OF LIGAND BINDING IN A BURIED NON-POLAR CAVITY OF T4 LYSOZYME: LINKAGE OF DYNAMICS AND STRUCTURAL PLASTICITY. Preprint at https://doi.org/10.2210/pdb182l/pdb (1995).
15. Souza, P. C. T. *et al.* Martini 3: a general purpose force field for coarse-grained molecular dynamics. *Nat. Methods* **18**, 382–388 (2021).
16. Souza, P. C. T. & Marrink, S. J. Martini 3 open-beta version. http://cgmartini.nl/index.php/martini3beta.
17. Wassenaar, T. A., Ingólfsson, H. I., Böckmann, R. A., Tieleman, D. P. & Marrink, S. J. Computational Lipidomics with insane: A Versatile Tool for Generating Custom Membranes for Molecular Simulations. *J. Chem. Theory Comput.* **11**, 2144–2155 (2015).
18. Martínez, L. Automatic identification of mobile and rigid substructures in molecular dynamics simulations and fractional structural fluctuation analysis. *PLoS One* **10**, e0119264 (2015).
19. Souza, P. C. T., Thallmair, S., Marrink, S. J. & Mera-Adasme, R. An Allosteric Pathway in Copper, Zinc Superoxide Dismutase Unravels the Molecular Mechanism of the G93A Amyotrophic Lateral Sclerosis-Linked Mutation. *J. Phys. Chem. Lett.* **10**, 7740–7744 (2019).
20. Strange, R. W. *et al.* Variable metallation of human superoxide dismutase: atomic resolution crystal structures of Cu-Zn, Zn-Zn and as-isolated wild-type enzymes. *J. Mol. Biol.* **356**, 1152–1162 (2006).
21. Salama-Alber, O. *et al.* Atypical cohesin-dockerin complex responsible for cell surface attachment of cellulosomal components: binding fidelity, promiscuity, and structural buttresses. *J. Biol. Chem.* **288**, 16827–16838 (2013).
22. Poma, A. B., Cieplak, M. & Theodorakis, P. E. Combining the MARTINI and Structure-Based Coarse-Grained Approaches for the Molecular Dynamics Studies of Conformational Transitions in Proteins. *J. Chem. Theory Comput.* **13**, 1366–1374 (2017).
23. Bernardi, R. C. *et al.* Mechanisms of Nanonewton Mechanostability in a Protein Complex

Revealed by Molecular Dynamics Simulations and Single-Molecule Force Spectroscopy. *Journal of the American Chemical Society* vol. 141 14752–14763 Preprint at https://doi.org/10.1021/jacs.9b06776 (2019).

46. Xiong, J. *et al.* Multi-instance learning of graph neural networks for aqueous pKa prediction.

*Bioinformatics* **38**, 792–798 (2022).

47. Dignon, G. L., Zheng, W., Kim, Y. C., Best, R. B. & Mittal, J. Sequence determinants of protein phase behavior from a coarse-grained model. *PLoS Comput. Biol.* **14**, e1005941 (2018).
48. Spinti, J. K., Neiva Nunes, F. & Melo, M. N. Room for improvement in the initial martini 3 parameterization of peptide interactions. *Chem. Phys. Lett.* **819**, 140436 (2023).
49. Hanwell, M. D. *et al.* Avogadro: an advanced semantic chemical editor, visualization, and analysis platform. *J. Cheminform.* **4**, 17 (2012).
50. Kim, T. & Im, W. Revisiting hydrophobic mismatch with free energy simulation studies of transmembrane helix tilt and rotation. *Biophys. J.* **99**, 175–183 (2010).
51. PyMOL. http://www.pymol.org/pymol.
52. Alessandri, R. *et al.* Pitfalls of the Martini Model. *J. Chem. Theory Comput.* **15**, 5448–5460 (2019).
53. Bayer, E. A., Lamed, R., White, B. A. & Flint, H. J. From cellulosomes to cellulosomics. *Chem. Rec.* **8**, 364–377 (2008).
54. Schoeler, C. *et al.* Ultrastable cellulosome-adhesion complex tightens under load. *Nat. Commun.* **5**, 5635 (2014).
55. Tsanai, M., Frederix, P. W. J. M., Schroer, C. F. E., Souza, P. C. T. & Marrink, S. J. Coacervate formation studied by explicit solvent coarse-grain molecular dynamics with the Martini model. *Chem. Sci.* **12**, 8521–8530 (2021).
